# HSSM: A Widely Applicable Toolbox for Hierarchical Bayesian Neurocognitive Modeling

**DOI:** 10.64898/2026.06.05.730398

**Authors:** Alexander Fengler, Yang Xu, Krishn Bera, Carlos Paniagua, Aisulu Omar, Michael J. Frank

**Author notes:** **For correspondence:**, (Alexander Fengler, Michael J. Frank). Department of Cognitive and Psychological Sciences, Brown University, Providence, RI, USA.

## Abstract

Computational models are central to cognitive neuroscience, but their rigorous application to experimental datasets is often constrained to a narrow set of canonical models that afford tractable analytical computations. We introduce the `HSSM` (Hierarchical Sequential Sampling Model) ecosystem, a Python toolbox that democratizes access to a broad, extensible array of neurocognitive process models through hierarchical Bayesian inference. Naturally leveraging simulation-based inference via likelihood surrogates, `HSSM` enables fast parameter estimation for models lacking closed-form likelihoods. Built atop PyMC and Bambi, `HSSM` provides a user-friendly formula syntax for specifying hierarchical mixed-effects regressions on model parameters, incorporating trial-by-trial neural or physiological covariates. The ecosystem allows fast model simulation and training data generation, as well as the neural network training utilities to deploy surrogate likelihood networks via HuggingFace. Contributions are designed to benefit not only the single researcher working on a problem, but organically, the entire research community. Together, the tools in the `HSSM` ecosystem bridge the interests of computational theorists as well as experimentalists, accelerating the cycle from model development to rigorous empirical testing.

## Introduction

Computational modeling has become pivotal to research in cognitive neuroscience and is increasingly used to lend insights into mechanisms of mental illness (***Huys et al., 2016; Friston et al., 2014; Wiecki et al., 2014; Huys et al., 2021; Maia and Frank, 2011; Halassa et al., 2025***). For example, the ability to accumulate evidence and then balance the rewards and costs associated with each choice is key to successful action selection. Failure in brain circuits governing decision-making, learning, memory, and cognitive processes can lead to maladaptive behaviors, which characterize many psychiatric disorders. Neurocomputational models, spanning implementational and algorithmic levels, have identified distinct neural mechanisms for a variety of cognitive functions, affording tractable quantitative fits to brain/behavior relationships (***Friston et al., 2014; Huys et al., 2016; Halassa et al., 2025***). By focusing on converging theoretical and cognitive neuroscience evidence, this approach offers a quantitative and computational interpretation of qualitative diagnostic features. There are two critical steps to this process: i) linking neurocognitive dynamic processes to formal models that can be quantitatively tested against experimental data; and ii) rigorous hierarchical Bayesian inference methods allowing researchers to estimate the parameters of these models (and their uncertainty), including how they vary with neural activity and across patient populations. Superior diagnostic classification can be achieved using such methods compared to purely data-driven approaches, including in patient populations with psychiatric or neurological disturbances (***Geana et al., 2022; Wiecki et al., 2014; Pedersen and Frank, 2020; Wiecki et al., 2016; Pagnier et al., 2024; Ging-Jehli et al., 2024***). The resulting parameter differences lend insight into brain mechanisms that dissociate forms disorders of memory, impulsivity, compulsivity, social decision making, and depression, amongst others (***Lawlor et al., 2020; Banca et al., 2016***).

As the field matures, it is increasingly important to move beyond standard popular models (***Cisek et al., 2009; Hawkins et al., 2015; Voss et al., 2019; Wieschen et al., 2020; Trueblood et al., 2021; Fengler et al., 2022; Rasanan et al., 2024b***,a) toward those that faithfully represent underlying cognitive and neural processes. There are, however, two core problems that prevent the broader research community from capitalizing on the full arsenal of theoretically meaningful models. First, the landscape of software toolkits available for efficient fitting of neurocognitive models is scattered. The overhead of working across disparate code bases, with little connective tissue, often limits experimental scientists to relying on a set of popular, canonical computational models and domain-specific toolboxes for testing against their experimental data. Second, many models that are theoretically meaningful and motivated by neural and cognitive principles are technically difficult to fit to experimental data. This crucial challenge stems from the lack of easily accessible closed-form analytic likelihood functions for these models, which are needed for fast Bayesian inference algorithms (***Cranmer et al., 2020; Radev et al., 2020; Fengler et al., 2021; Navarro and Fuss, 2009***). In particular, while it is often trivial to specify a simulator for a given generative model (going from model and “parameters to data”), the inverse problem is not so simple. When interrogating a given experimental dataset, the challenge is to go the other way around: to quickly infer which model and under which parameter configuration would have most likely given rise to that pattern of data. Solving this inverse problem can be surprisingly intractable even when forward simulation is straightforward. Box 1 provides a concrete illustration.

As a result, the vast majority of researchers limit themselves to a small class of canonical models with accessible likelihood functions for analytical convenience (***Ratcliff, 2006; Brown and Heathcote, 2008***). Because these models might not be sufficient to account for the underlying processes, this shortcut can result in misleading interpretations of model results. Indeed, our predecessor toolbox, `HDDM` (***Wiecki et al., 2014***), collected more than a thousand citations through applications, while limited to a single canonical cognitive process model, the DDM (***Ratcliff, 2006***). For many of these applications it is likely that alternative models may have been the more realistic, but were not considered due to the simple lack of conveniently accessible software infrastructure. While we eventually extended the `HDDM` toolbox (***Fengler et al., 2022***) in recognition of our methodological developments (***Fengler et al., 2021***), our work here in turn recognizes that `HDDM` was never fundamentally designed for the model-level flexibility that was unlocked by the contemporary computational tools.

Recent methodological advances apply deep learning in combination with simulation-based inference (***Cranmer et al., 2020; Radev et al., 2020; Fengler et al., 2021; Miller et al., 2022; Radev et al., 2023; Huang et al., 2026***) to enable inference over a much larger spectrum of models, without requiring analytical likelihoods. The corresponding approaches require access to only a simulator for training a neural network that can learn the likelihoods (a likelihood approximation network or LAN; Fig 2), which can then be used for fast Bayesian inference when fitting experimental data. (We note that there also exist simulation-based inference methods for directly approximating posterior distributions (***Radev et al., 2020; Tejero-Cantero et al., 2020***). While such approaches can be powerful, to date they do not lend themselves naturally to arbitrary hierarchical designs or flexible estimation of how model parameters vary by neural activity etc.)

**Figure 1.**
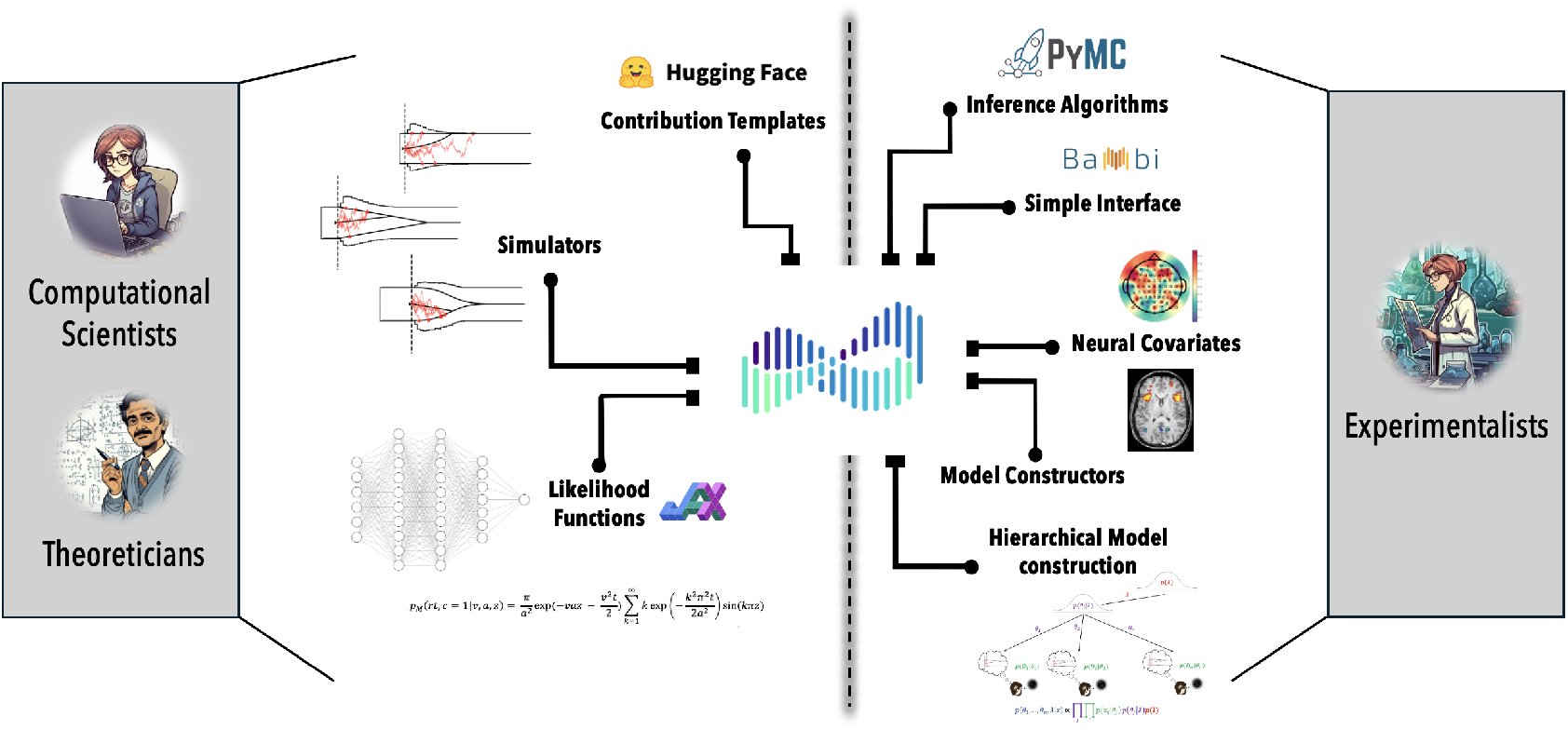
Basic Conceptual overview of the `HSSM` Ecosystem. HSSM provides a platform for theoreticians and computational scientists to contribute (and stress test) new models and for experimentalists to test their empirical data against a larger, expanding array of such models. The resulting cycle contributes to accelerate translation between theoretical and experimental contributions to the field of computational cognitive neuroscience.

**Figure 2.**
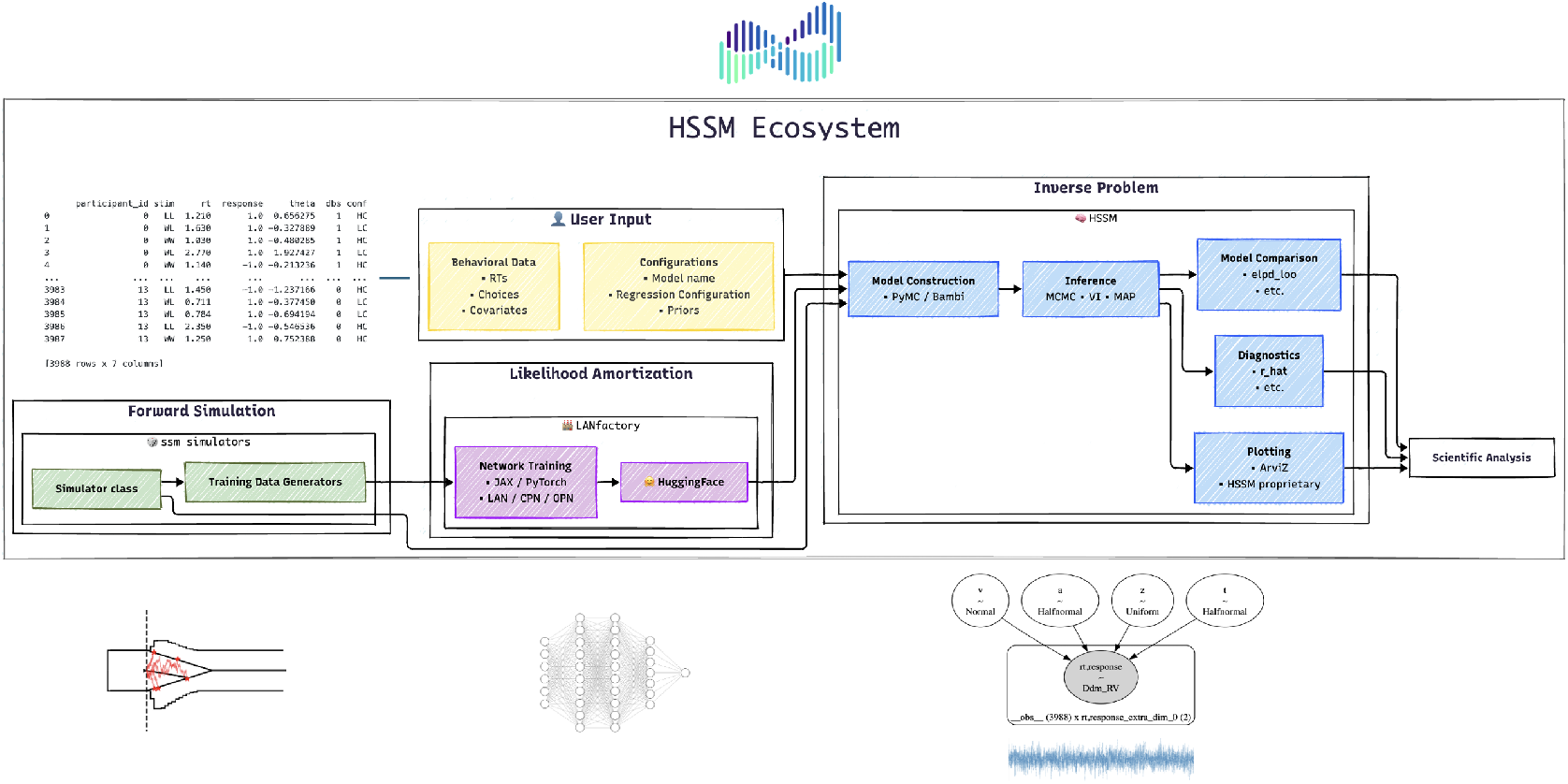
Overview of the HSSM ecosystem. The core ecosystem consists of three interoperable Python packages (left column) that span the end-to-end workflow from cognitive model simulation to scientific analysis via Bayesian parameter inference (right panel). `ssm-simulators` provides fast forward simulators for a broad, extensible set of cognitive process models, and produces the training data required for likelihood surrogate approximation. `LANFactory` trains lightweight neural networks — likelihood approximation networks (LANs), choice probability networks (CPNs), and omission probability networks (OPNs) — using JAX or PyTorch, and exports them as ONNX files that can be uploaded to a shared HuggingFace repository for community reuse. `HSSM` is the user-facing hub: it retrieves the relevant simulator and likelihood, combines them with user-supplied behavioral data and a formulaic model description, constructs the model via PyMC and Bambi, and supports full Bayesian inference (MCMC, VI, MAP) alongside posterior predictive plotting, diagnostics, and model comparison. The right panel traces the full pipeline from forward simulation through likelihood amortization through to the inverse problem of parameter inference and downstream scientific analysis.

To unlock the inherent potential to impact scientific workflows more broadly, the `HSSM` ecosystem provides a collection of toolboxes that (i) allow computational experts to contribute new models and methods (developer-API), and (ii) provides a user-friendly interface for experimental scientists to rigorously interrogate their data (user-API). This allows researchers to not only fit a broad array of models, but to easily construct statistical models to assess how their parameters may vary with neural or other physiological activity, or whether they differ between healthy and patient populations. It also provides functions that allow users to perform various model validation checks, such as posterior predictive checks, dynamic cartoon plots of stochastic model behavior, parameter recovery, and model selection.

### Box 1.

Illustration of the forward-inverse asymmetry with a basic statistical example.

For intuition, consider a Bernoulli model that generates binary outcomes such as coin flips with some probability *θ*. It is straightforward to generate a dataset by randomly generating 1’s and 0’s according to *θ*. This is the *forward process*,

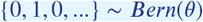

As a researcher we are however usually confronted with the problem of *inferring p* after observing a random sequences of 1’s and 0’s. The usual Bayesian approach to this problem makes use of the proportionality (Bayes’ Rule),

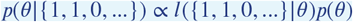

where *p*(*θ*|*data*) or *p*(*θ*|{1, 1, 0, …}) is the *posterior, l*({1, 1, 0, …}|*θ*) is the *likelihood* and *p*(*θ*) is a suitable *prior*.

Notice that the process to generate (simulate) data via *Bern*(*θ*) (you could be physically throwing coins) can be totally different from the computations you want to carry out when evaluating the likelihood of some observed sequence following this process. Deriving the likelihood may seem easy here (*θ*^#1’s^ ∗ (1 − *θ*)^#0’s^), however in general it may be mathematically tedious, hard or even impossible, even for models that can seem suspiciously easy to generate data from. However for any traditional approach to Bayesian inference, it is crucial to have the tractable likelihood term, otherwise we can not compute the right hand side of Bayes’ rule. Such a tractable function has been derived and put to good use for some cognitive process models, such as the drift diffusion model (DDM; ***Navarro and Fuss (2009***)), however even minor variations of these drift diffusion models (which may be more informed by neural dynamics, for example) render likelihood derivations and/or computations intractable or at least computationally much more demanding. Indeed, the space of cognitive models with tractable analytic solutions is far smaller than the space of popular or theoretically interesting models(***Cisek et al., 2009; Hawkins et al., 2015; Voss et al., 2019; Wieschen et al., 2020; Trueblood et al., 2021; Fengler et al., 2021; Rasanan et al., 2024b***,a). Researchers have traditionally required computationally very expensive Monte Carlo simulation methods for parameter fitting (***Beaumont et al., 2002; Beaumont, 2010; Wood, 2010; Turner and Van Zandt, 2012; Turner and Sederberg, 2014; Turner and Van Zandt, 2018; Huys et al., 2016***). For many concrete use-cases — e.g. testing model variants such as how parameters may vary by task condition, neural activity, or hierarchically across many participants — such methods can take weeks for any individual model (***Radev et al., 2020; Tejero-Cantero et al., 2020***).

We hope to incentivize each community to interact with the `HSSM` ecosystem for the benefits specifically tailored to their needs. Computational experts may benefit from a larger audience of prospective users for testing their theoretical ideas. Access to a very broad set of cognitive models and the ability to generate very flexible hierarchical models around such a model core, in turn, helps experimentalists test manifold specific theories against their empirical data.

We will begin with an overview of the core contributions of our `HSSM` ecosystem. We then provide a package-wise overview of the capabilities and contextualize design choices. Accordingly, we will first explain the vision and principles behind the overall development and how the packages in the ecosystem interact and support each other. Second, we will go through the core capabilities of each, the `HSSM`, `ssm-simulators` and the `LANFactory` packages in turn. We will then discuss related work and how `HSSM` fits into the overall landscape of software to support cognitive process modeling. Finally, before we conclude, we will discuss planned future developments and our approach to long-term support for the collection of tools presented in this paper. This long term support has in mind developments to embrace a wide variety of emerging agentic workflows, allowing scientists to achieve more without manual painstaking step-throughs of a pipeline for each alternative model, as well as lower the barrier of entry for beginners from dataset to hierarchical cognitive process models.

## HSSM Ecosystem: Overview

At the core of our software ecosystem is the Hierarchical Sequential Sampling Modeling (HSSM) Python package, which acts as the hub through which researchers can flexibly build and perform parameter inference on complex hierarchical models tailored to the specifics of their experimental datasets. The package is general and in principle supports *any* generative model of behavior as well as how dynamic brain processes can dynamically alter these generative processes within and across trials and/or experiment conditions. As an initial test bed we have extended functionality for a wide range of SSMs and reinforcement learning models, including any modular combination of the two (see the dedicated section below). Without loss of generality we also will extend the package to include models of continuous reports (e.g., perceptual and working memory judgments, continuous psychophysics, motor control and more) (***Burge and Bonnen, 2025; Rasanan et al., 2024a***).

Leveraging recent deep learning based developments in simulation based inference (***Cranmer et al., 2020; Radev et al., 2020; Fengler et al., 2021***), `HSSM` provides a simple model-building interface with which users can easily construct hierarchical Bayesian models upon an expanding and user-extensible list of neurocognitive process models. `HSSM` allows everything from standard analytical closed-form likelihoods (***Brown and Heathcote, 2008; Navarro and Fuss, 2009***) when available, to any kind of likelihood approximation (***Fengler et al., 2021; Radev et al., 2023; Boelts et al., 2022; Miller et al., 2022***), including but not limited to those we have developed and validated (***Fengler et al., 2021, 2022***).

While we allow a range of algorithms to fit parameters, a core focus of HSSM is on Hierarchical Bayesian estimation. This statistical approach optimizes the tradeoff between random and fixed effects models, capitalizing on the complete dataset by naturally pooling information from the group to inform estimation of individual subject parameters. Thus, the posterior distribution of individual (*θ*) and group (*λ*) level parameters can be inferred from the data **x** as:

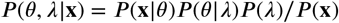

The method capitalizes on statistical strength shared across individuals, exploiting the extent to which subjects are similar to each other, and optimizes the tradeoff between random and fixed effects to improve parameter estimation. Model variants can be tailored to specific hypotheses (e.g., effect of reward incentives or trial-wise brain activity on a given parameter), and to test how model parameters may vary within- or between individuals e.g., due to stress, symptoms, brain stimulation (***Ging-Jehli et al., 2024; Pedersen et al., 2021; Kuhn et al., 2025; Pagnier et al., 2024***).

Crucially, the abstract design of HSSM, in principle, allows essentially any cognitive model (and beyond) to be incorporated here. We provide a range of models with `HSSM`from the outset, including a wide variety of evidence accumulation models (DDM, LBA, race models with multiple choice options, conflict models, etc), as well as interface for models that evolve over experience, such as reinforcement learning models, as well as any combination of the two types of within- and across-dynamics (see below). However, this flexibility is merely a sample of the expansion that our framework allows.

The process of model building and estimation is seamless: once the user specifies a model, the package selects the appropriate likelihood function — whether analytic, neural network based, or some other surrogate — and users can then perform fully hierarchical Bayesian inference using state-of-the-art Markov Chain Monte Carlo methods, including gradient-based samplers and GPU acceleration, yielding speeds that are orders of magnitude faster than traditional simulation-based approaches (***Turner and Sederberg, 2014; Holmes, 2015***). The main power of `HSSM` lies in the ease with which these interpretable, low-dimensional computational models can be combined with any trial-by-trial covariate (fMRI, EEG, spikes, eye-tracking, skin-conductance, etc.), using the broadly popular Python programming language. This promises to substantially enhance discovery of links between brain and cognitive dynamics across a wide range of phenomena, as well as how these are altered in disorders of brain function. Through HSSM, experimentalists can access this large bank of cognitive process models and benefit directly from innovations in likelihood computation without requiring technical sophistication.

Conversely, `HSSM` serves as the front-end through which theoreticians and computational scientists can make new models available to the community and benefit from wider adoption through a trusted ecosystem which also provides tools for model validation. This connective tissue will fundamentally enable a new paradigm for how computational and experimental scientists interact, unlocking great potential for significant speedup of research cycles.

To support a prototypical and convenient end-to-end workflow for computational scientists, our ecosystem contains two auxiliary packages. The `ssm-simulator` package is built to support fast simulation for a broad, extensible class of cognitive process models. It integrates seamlessly with the second auxiliary package, `LANFactory`, which allows users to train artificial neural networks in the service of likelihood evaluation (***Fengler et al., 2021, 2022***). `LANFactory`, in turn, integrates seamlessly with `HSSM`: once a likelihood network is trained, it is made available for all users to test the corresponding model against their data across any inference setting, without requiring more computationally costly simulations or network training. All tools are constructed with the following principle in mind to motivate adoption: be maximally helpful for beginner users, and minimally intrusive on workflow choices for expert users. For example, users can contribute any other likelihood evaluator to `HSSM` even if it is not trained through `LANFactory`. We showcase examples of such a workflow via likelihood ratio estimators extracted form the `BayesFlow` python library (***Radev et al., 2020***). More on the core development principles behind our ecosystem below. Lastly, as we will show later, `HSSM` itself has developer and advanced APIs to allow people to siphon off useful components and use them directly through low-level probabilistic programming libraries such as PyMC (***Abril-Pla et al., 2023***).

### Components and Interplay

The `HSSM` ecosystem consists of three underlying Python packages, designed for maximum interoperability but without sacrificing the ability to use any of them independently. If a user would like to simply have access to a fast and flexible simulation package, `ssm-simulators` has independent utility, without necessarily tying it to probabilistic modeling in `HSSM`. Conversely, `HSSM` users can fit models to their data and run prior and posterior predictive checks without needing to concern themselves with such functions leverage the `ssm-simulators`. No aspect of `HSSM` forces a user into explicit knowledge around it’s simulation backend. However if users are interested in the intersections, the modularity of the ecosystem serves as a powerful engine for contribution and creative usage of the components.

1. **ssm-simulators**. The `ssm-simulators` package provides fast trajectory simulation from a large variety of SSMs (including any new user-defined custom models). This allows, (i) visualization of model behavior, (ii) prior and posterior predictive checks and (iii) the training data foundation for downstream training of likelihood surrogates (amongst other things) (***Fengler et al., 2021; Leng et al., 2024***).
2. **LANFactory**. The lightweight `LANFactory` package interfaces naturally with the structure of training data coming out of `ssm-simulators` and is used to train LANs (and the related CPNs and OPNs; see text and details in the section dedicated to the `LANFactory package`). These networks can then be converted into ONNX format and uploaded to HuggingFace for the larger community to benefit from within and outside of HSSM.
3. **HSSM**. The `HSSM` package is the heart of the ecosystem, built around PyMC (***Abril-Pla et al., 2023***), enabling statistical workflows around Hierarchical Bayesian Inference and model validation. Users can access a range of underlying cognitive process models (including a wide variety of sequential sampling models) and construct parameter-wise hierarchical mixed-effect regressions with full freedom on the specification of priors. Model construction is facilitated via the Bambi (***Capretto et al., 2022***) library. The `ssm-simulators` package provides the forward simulators, and the `LANFactory` package provides the surrogate Likelihoods for analytically intractable models. `HSSM` combines these pieces flexibly.

The packages can be used together, however they also serve as independent end-points. A user may want to access the fast simulators in the `ssm-simulators` package, without worrying about either likelihood approximation nor any downstream parameter inference. Likewise, a user may want to train a LAN on a model that `ssm-simulators` fundamentally doesn’t support. Lastly, and we recognize this as the most likely case, an experimentalist may want to use `HSSM` in isolation, without concerning themselves explicitly with network training nor any user-facing simulation code logic. For this last user, `ssm-simulators` and `LANFactory` serve as solid (but not necessary) bedrock for model contributions to the ecosystem, not as independently used entities. Figure 2 provides a conceptual breakdown of the ecosystem design.

Our packages are unified by the following overall design principles:

1. **Flexibility**. *High ceiling, low floor*. We provide guidance without strict guardrails. We develop key abstractions to be helpful for the beginner user who wants the shortest path to empirical data analysis, as well as low-level access points for the advanced user who want full controllability to use bits and pieces of the ecosystem inside their custom workflows.
2. **Extensibility**. *Designed for community contribution*. Our abstractions prioritize user extensibility, to enable community contributions with minimal intervention points.
3. **Innovation Inheritance**. *Benefit from and support the broader ecosystem*. We designed the ecosystem so that it can benefit from continual innovations happening in the network of upstream dependencies, including probabilistic programming and simulation based inference.

We will return to these principles and extensions in the discussion section. In what follows, we first provide a detailed breakdown of each of the basic packages in the HSSM-ecosystem.

## HSSM

### Place in the Ecosystem

HSSM is the user-facing hub of the ecosystem, where simulation and likelihood approximation converge to enable principled statistical inference. It draws on simulators registered in `ssm-simulators` for forward modeling, analytical or surrogate likelihoods trained via `LANFactory` (or supplied by other means) for parameter estimation. It seamlessly combines these components transparently behind a high-level model-building interface. For the majority of end-users — experimentalists fitting models to behavioral and neural data — `HSSM` is the only package they need to interact with directly. At the same time,`HSSM` is designed to be a component in broader workflows rather than a closed system. Its low-level API exposes reusable PyMC distributions, and its compiled log-probability functions can be extracted and passed to external sampling libraries. This positions `HSSM` not only as the downstream consumer of the ecosystem’s other packages, but also as a flexible model-construction engine that researchers can integrate into their own computational pipelines. Figure 3 provides a conceptual breakdown of the package design.

**Figure 3.**
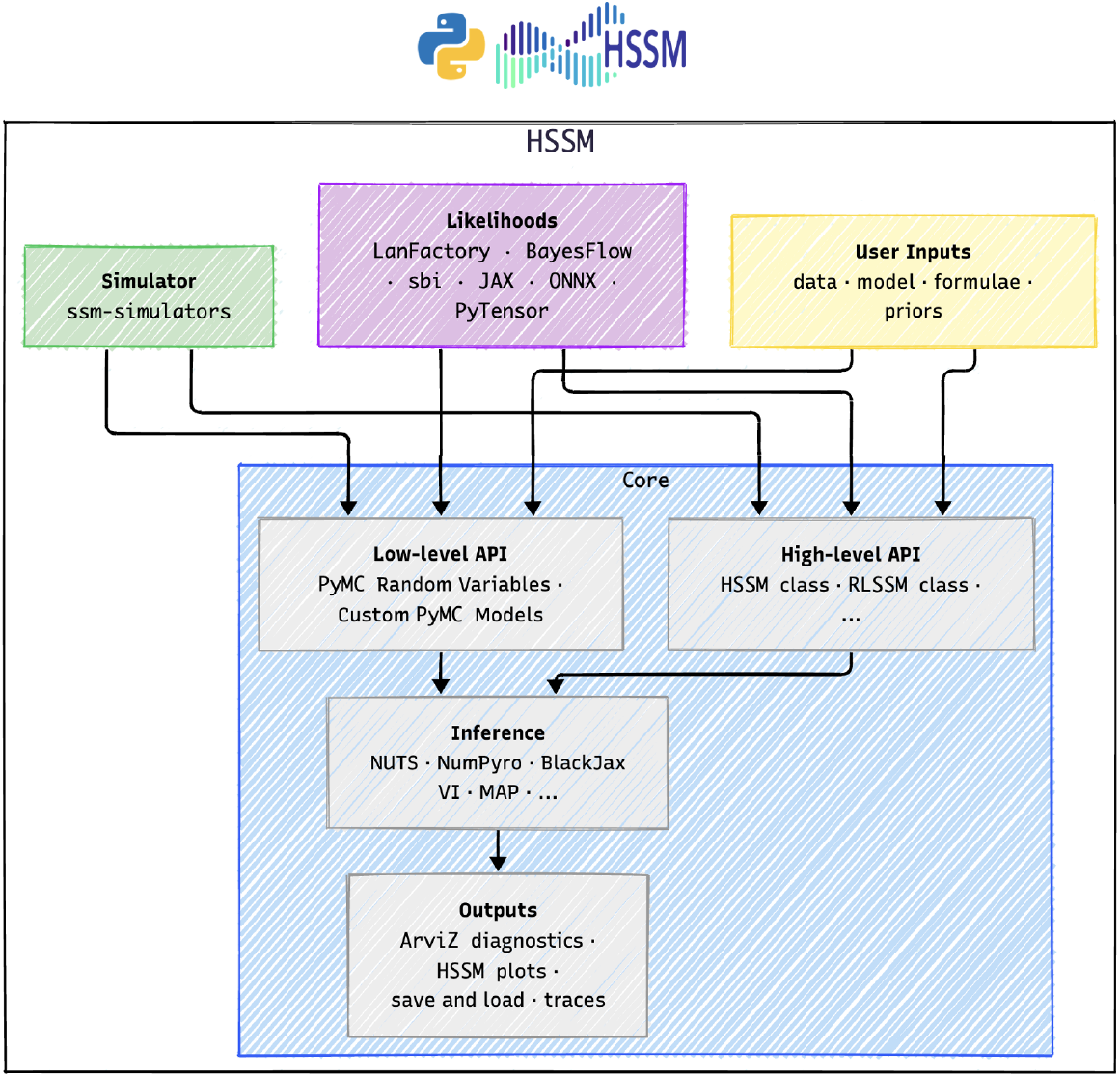
Architecture of the HSSM package. `HSSM` is designed around two complementary APIs at its core: the *high-level API* (`HSSM` class, `RLSSM` class, etc.) constructs hierarchical Bayesian models automatically from user-supplied data, model choice, formulae, and priors; the *low-level API* exposes pre-assembled PyMC random variables and supports embedding HSSM likelihoods in entirely custom PyMC models. The likelihood layer accommodates multiple sources, including surrogates produced via `LANFactory`, `BayesFlow`, or `sbi` and deployed through JAX, ONNX, or PyTensor. Forward simulators are drawn from `ssm-simulators` for prior and posterior predictive sampling; users can supply their own. The assembled model is a standard PyMC distribution and can be fit with a wide range of inference backends — gradient-based MCMC via NUTS, NumPyro, or BlackJax; variational inference; MAP estimation; or external samplers. Outputs are compatible with ArviZ diagnostics and HSSM’s plotting utilities, and models can be persisted via native save/load methods. See Figure 3—figure supplement 1 for the full internal architecture. **Figure 3—figure supplement 1**. HSSM Conceptual Architecture: Detailed

### Core Functionality

The `HSSM` package provides four interrelated capabilities that together support an end-to-end Bayesian modeling workflow for cognitive process models.

First, `HSSM` offers a high-level model construction interface through which users specify a cognitive process model, define hierarchical mixed-effects regression structures on its parameters, and assign prior distributions — all via a concise formula syntax inherited from Bambi. This allows users to express complex experimental hypotheses (e.g., that drift rate or learning rate varies by condition and is modulated by trial-wise neural activity, with random intercepts across participants) in just a few lines of code.

Second, `HSSM` provides likelihood source management. For each registered model, the package helps with the selection of the appropriate likelihood function — whether an analytical closed-form expression, a LAN-based neural network surrogate, or a user-supplied custom function. Beginner users can rely on defaults and interacts with a uniform interface regardless of the underlying computational machinery.

Third, through PyMC (***Abril-Pla et al., 2023***) `HSSM` supports multiple inference backends. Users can perform fully hierarchical Bayesian inference via Markov Chain Monte Carlo sampling (including gradient-based samplers such as NUTS), variational inference, or optimization-based point estimation, all through a consistent API. GPU acceleration is available for compatible backends, which, in particular, the neural network based likelihoods can use to great effect, yielding inference speeds that can be orders of magnitude faster than traditional simulation-based approaches.

Fourth, `HSSM` includes a suite of model validation and visualization tools. These include posterior and prior predictive checks, quantile probability plots for assessing fit to choice-RT distributions, and model cartoon plots that overlay inferred parameters onto visual representations of the generative process. Together with native integration with ArviZ for standard diagnostics (trace plots, summary statistics, model comparison via WAIC and LOO), these tools support rigorous assessment of model adequacy.

### Example: Loading Data

#### Box 2.

The `cavanagh_theta` dataset

**Box 2—figure 1.**
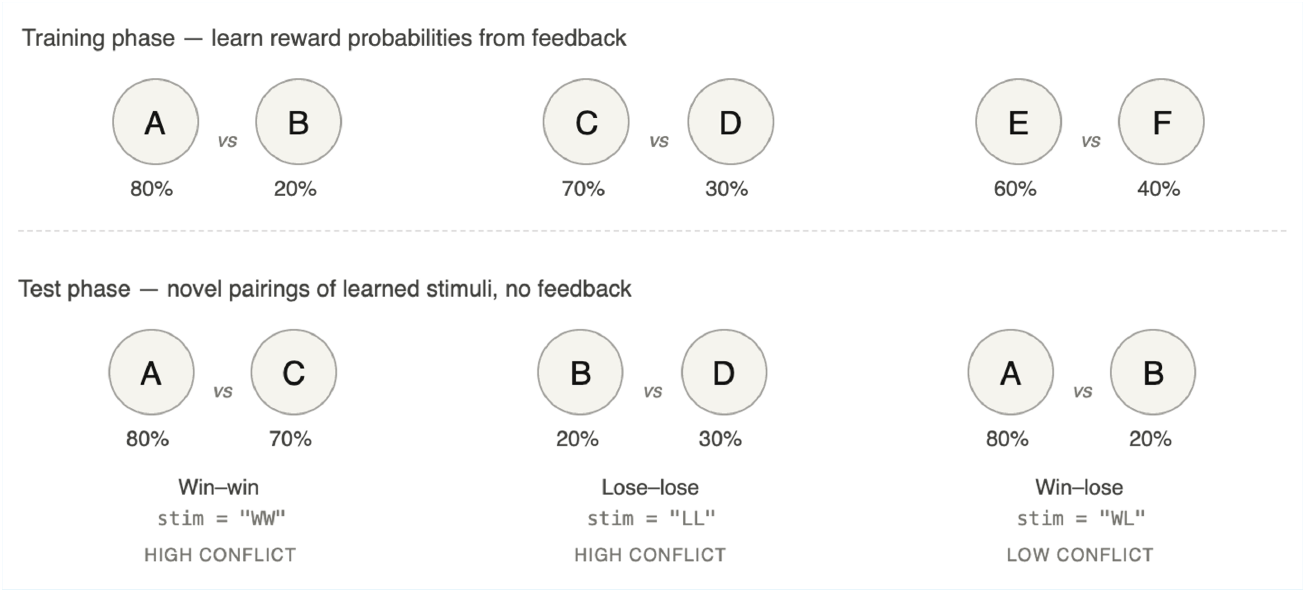
Simplified task structure for the “`cavanagh_theta`” dataset. Training pairs (top) and example test-phase pairings with corresponding `stim` values (bottom).

The example dataset used throughout this paper is derived from ***Cavanagh and Frank (2014***), who employed a probabilistic selection task (***Frank et al., 2004***). Participants first learned the reward probabilities of three stimulus pairs through trial-and-error feedback: AB (80%/20%), CD (70%/30%), and EF (60%/40%). In a subsequent test phase — the portion used here — they chose between all possible pairings without further feedback, yielding three condition types: *win–win, lose–lose*, and *win–lose*. The columns *used in this paper* are `rt` (response time in seconds), `response` (binary choice), `stim` (trial condition: `WW`, `LL`, `WL`), `participant_id`, and `theta` (trial-wise frontal-midline EEG theta power, used here as a continuous neural covariate). We *note* that, occasionally, to simplify figures didactically, we collapse the two “high conflict” conditions in the `stim` column.

`HSSM` ships with an example dataset which we call “`cavanagh_theta`”. This dataset is inherited from the predecessor `HDDM` toolbox (***Wiecki et al., 2014***). See ***Cavanagh and Frank (2014***), for details on the underlying study from which the dataset derives, however for convenience, in Box 2 we explain the aspects that help the reader understand the examples in this paper.

Listing 1 shows how to load this dataset. We will utilize this data as a running example for the remainder of our hssm code-snippets, wherever possible. Occasionally, we will rely on simulated data.

**Listing 1.**
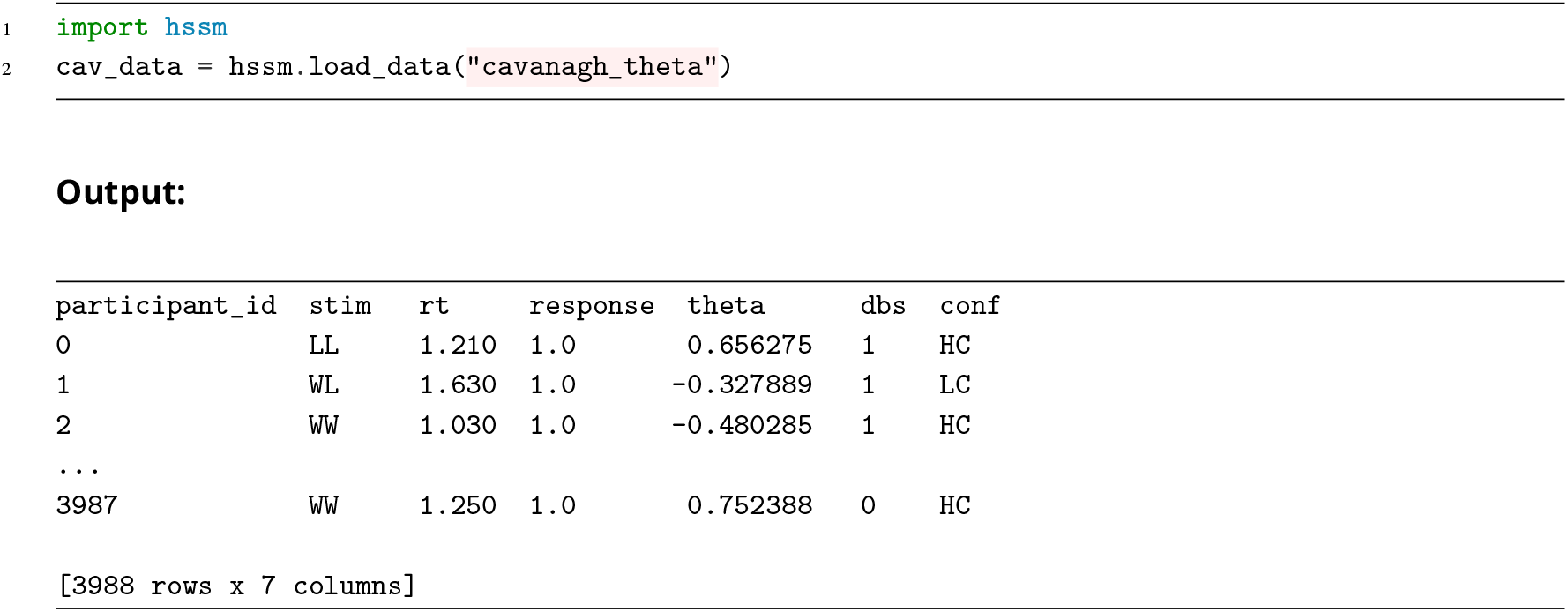
Loading a dataset that is included in the `HSSM` package.

**Listing 2.**
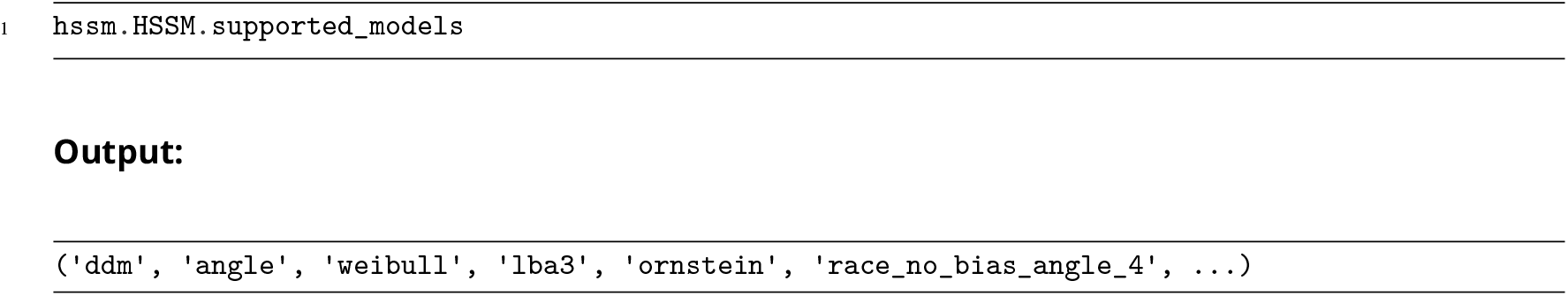
Getting the list of models currently supported in `HSSM`

`HSSM` can check for models that are *fully registered* in the ecosystem, i.e. those for which we have both a *simulator* and a valid *likelihood* available, providing all ingredients to construct PyMC random variables. A *fully registered* model implies that we will be able to do *parameter inference* as well as forward simulation, the latter facilitating *prior and posterior predictive sampling*. Listing 2 shows how to get the current list of fully supported models.

The `HSSM` documentation shows a variety of ways in which the package can handle partial construction (e.g., if we only provide a custom likelihood, ignoring the simulator during exploratory workflows). Users will be warned about the loss of particular levels of functionality when exploiting this flexibility.

### Example: Minimal HSSM Workflow

In this section we showcase the minimal version of what working with `HSSM` looks like in practice. Relying on defaults, we can accomplish a basic Bayesian workflow in very few lines of code. We reuse our “`cavanagh_theta`” dataset throughout.

First, we instantiate an `HSSM` model, passing the dataset and the type of cognitive process model we would like to test (here “`ddm`”). The print method provides rich textual output on all important aspects of the model: Parameters that are fit, priors for each, lapse probabilities as well as the chosen lapse distribution. The `.graph()` method allows us to leverage graphical model representations produced via PyMC (***Abril-Pla et al., 2023***).

To perform parameter inference via the standard PyMC NUTS sampler implementation and reasonable default settings you can use the simple, argument-less call “`hssm_model_simple.sample()`.

Once we have our posteriors, which we can access either via our returned, `idata_simple`, or by making use of the attached `.traces` object.

**Listing 3.**
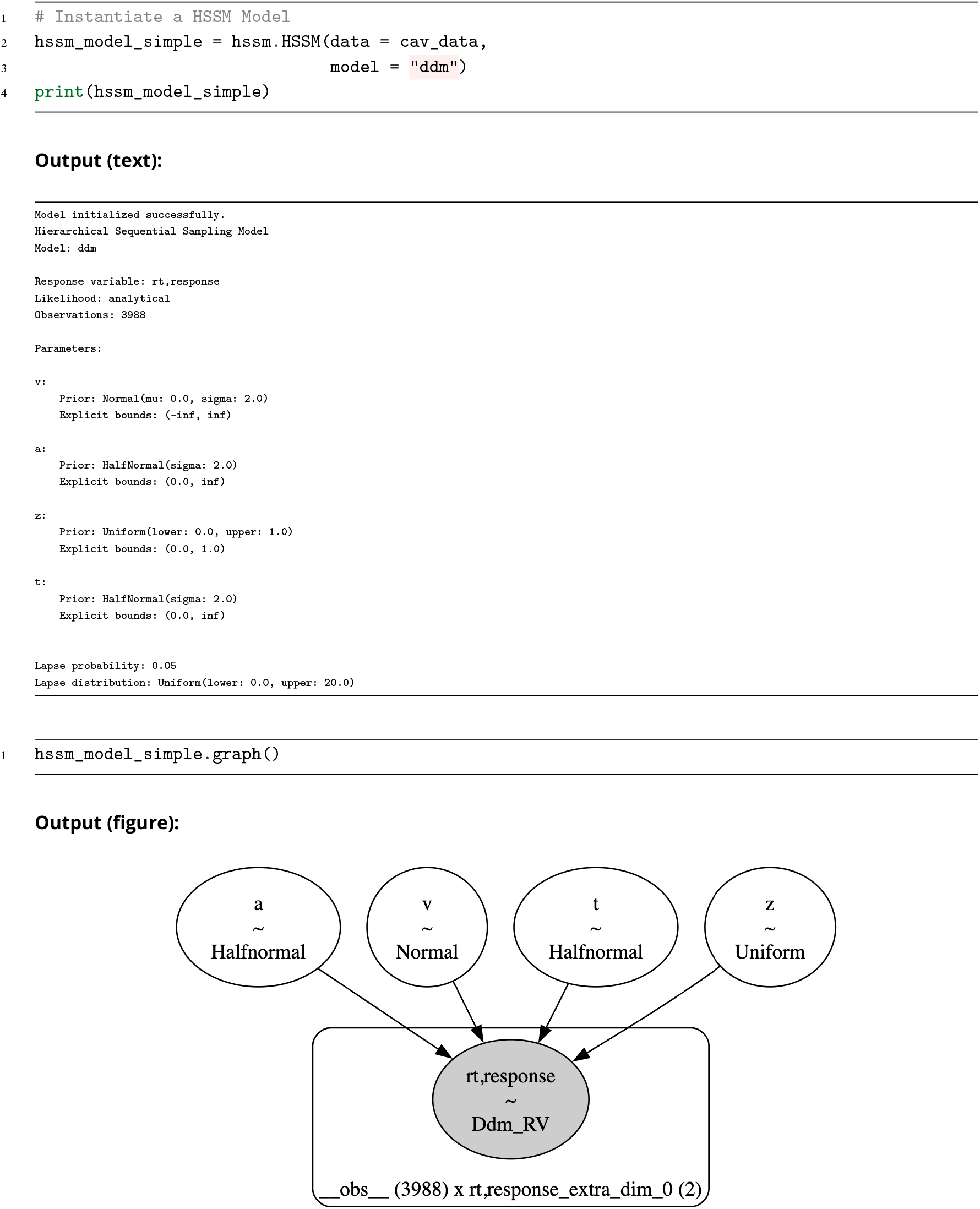
Illustrating a specified `HSSM` model. We provide a custom string for a simple readout of prior choices and the cognitive model parameters. Inheriting from the PyMC ecosystem, we can also get a simple graphical model depiction of our model via the `graph` method.

This is an `arviz.InferenceData` object, and we can now simply use it to access ArviZ (***Kumar et al., 2019***) functionality (e.g. the `plot_trace` function as illustrated).

### Specifying Regressions

`HSSM` allows you to specify arbitrary hierarchical mixed-effect regressions on each of the core parameters of our chosen cognitive process model. The Bambi (***Capretto et al., 2022***) backend constructs the parameter wise design matrices.

**Listing 4.**
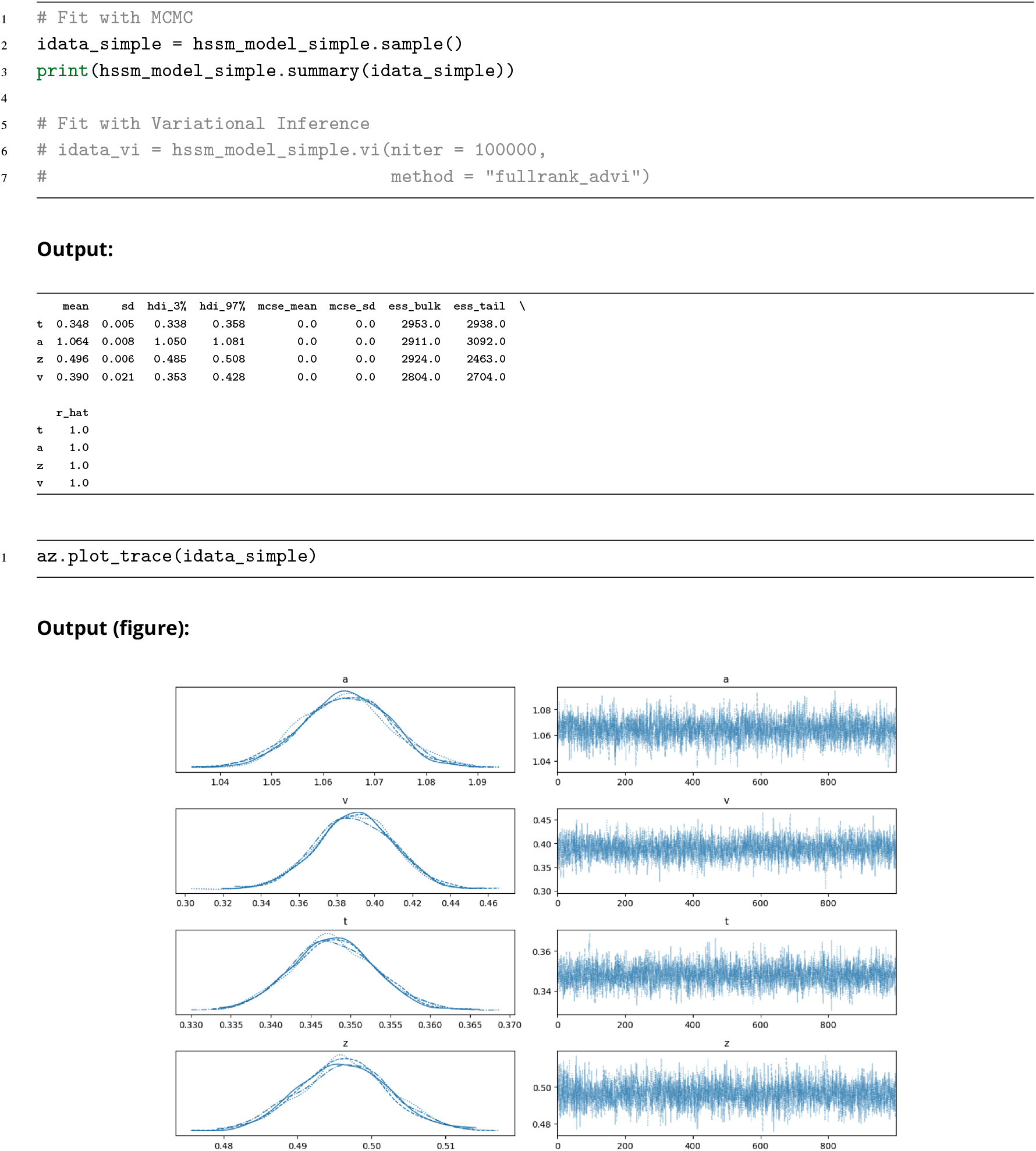
`HSSM` natively returns ArviZ `idata` objects which hold our traces from a model fit. We can then access the full plotting capability of ArviZ to illustrate results. We showcase the basic trace plot above.

In our example below, we chose the “`angle`” model (see Box 3), defaults for model parameters `t`, `z` and `theta` (the angle of the boundary collapse) and we add a regression to `a` and `v`.

In particular, we let the `a` parameter vary by our `stim` dimension (a categorical variable, referring to the specific type of visual stimuli, “low conflict” or “high conflict”), and we let the `v` parameter vary by `stim`, as well as a random intercept via the “`(1|participant_id)`” term.

Bambi allows us to change from centered to non-centered parameterizations (***Gelman, 2004; Betancourt and Girolami, 2015; Papaspiliopoulos et al., 2007***) via the `noncentered` argument.

The non-centered parameterization is often useful in the context of hierarchical models, however it is no cure-all and we suggest the user actively explore this setting and the scenarios in which it may or may not help with convergence. A full theoretical discussion is beyond the scope of this paper, however Box 4 provides a quick intuitive explanation. In future work we will allow this parameterization to target each parameter individually to provide greater freedom in dealing with parameterization related convergence issues.

#### Box 3.

Running example: The angle DDM

**Box 3—figure 1.**
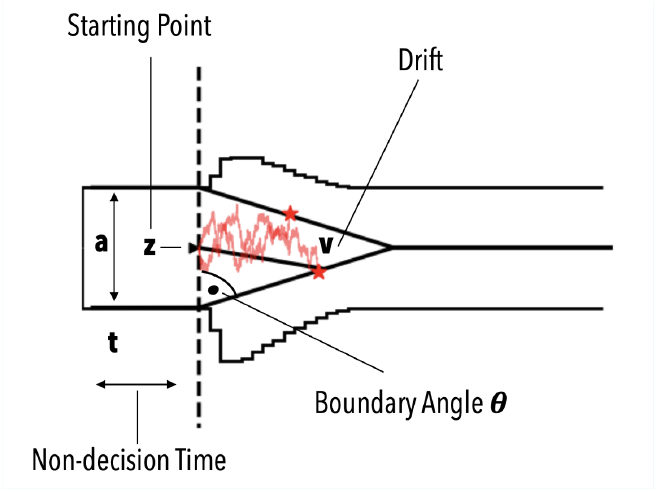

Schematic of the angle DDM.

The standard drift diffusion model (DDM) describes a two-alternative decision as a noisy accumulation of evidence toward one of two flat decision boundaries (***Ratcliff, 2006***). It is parameterized by a drift rate *v* (rate of evidence accumulation), a boundary separation *a* (distance between the two decision boundaries), a starting point *z* ∈ [0, 1] (relative bias toward one of the two boundaries), and a non-decision time *t* (time spent on processes other than evidence accumulation, e.g., perceptual encoding and motor execution). The decision variable *X*(*τ*) evolves over within-trial time *τ* according to:

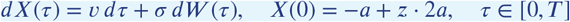

where *W* (*τ*) is a standard Wiener process and *σ* is a noise scale (typically fixed). A response is recorded when *X*(*τ*) first reaches −*a* (lower boundary, response = −1) or *a* (upper boundary, response = +1). The observed reaction time is *τ* + *t*, where *τ* is the first-passage time. For this model, an analytical likelihood is available (***Navarro and Fuss, 2009***) and forms the basis of much of the SSM literature. The *angle DDM* (***Fengler et al., 2021; Hawkins et al., 2015***) extends the standard DDM with a single additional parameter *θ* that determines the angle of linearly collapsing decision boundaries: the upper boundary becomes *a* − tan(*θ*) ⋅ *τ* and the lower boundary becomes −*a* + tan(*θ*) ⋅ *τ*. When *θ* = 0, the angle DDM reduces exactly to the standard DDM.

### Jointly modeling across- and within-trial decision dynamics: RL-SSMs

Recent work in computational cognitive science has increasingly focused on models that incorporate higher-order cognitive processes (e.g. learning, control, memory) into the trial-wise dynamics of decision making (***Pedersen et al., 2017; Miletić et al., 2021; Fontanesi et al., 2019; Ballard and McClure, 2019; Fengler et al., 2022***). In contrast with the conventional paradigms (such as perceptual discrimination and lexical decisions) that assume trial observations to be i.i.d., this class of models induces sequential dependencies across trials. This in turn enables researchers to formalize and test richer and more expressive theories of cognitive processes that modulate decision dynamics, including trial-and-error learning, load-dependent cognitive control, and memory-guided action selection (***Bera et al., 2025; McDougle and Collins, 2021***).

A canonical class of such models is Reinforcement Learning – Sequential Sampling Models (RLSSM). It combines the across-trial learning dynamics with the within-trial decision dynamics to jointly account for the choice and response time distributions (Figure 4). The core idea is that the learning process and the decision process are not independent but jointly generate the observed behavioral data. In the simplest version, RLSSMs replace the softmax action selection with an SSM, so that the trial-wise value estimates from the RL process feed forward as drift rates into the evidence accumulation process. This creates a generative model for the full joint distribution *p*(choice, RT ∣ parameters) on every trial, where the drift rates are non-stationary because they depend on the evolution of learned valuations.

**Figure 4.**
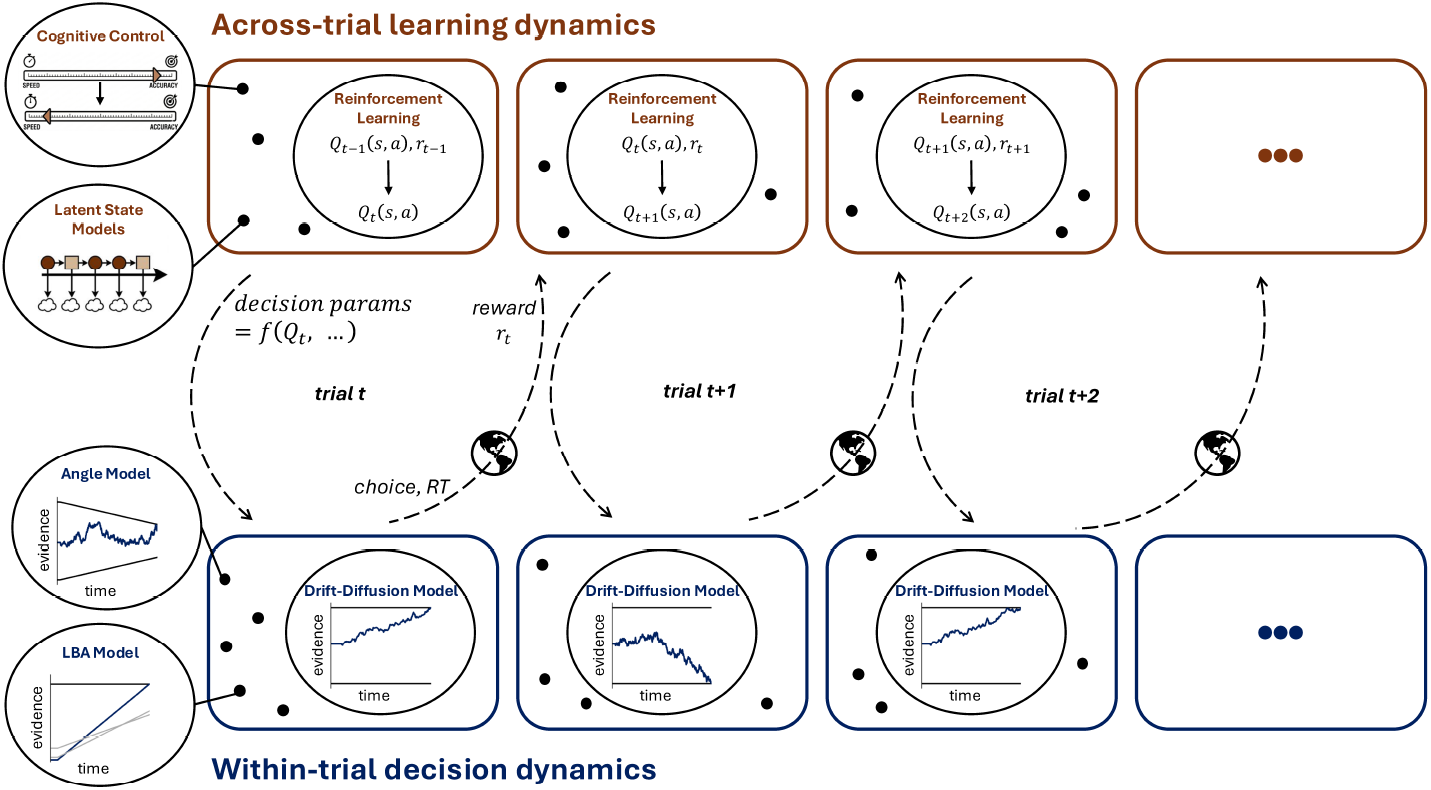
Reinforcement Learning - Sequential Sampling Models (RLSSM). RLSSM is a canonical example of combining across-trial cognitive processes such as reinforcement learning (RL) alongside within-trial decision processes such as DDMs and other SSMs. The DDM trial-by-trial parameters are modified via RL, e.g., to capture how the value of decision variables may evolve with experience or feedback in the task. The model produces a joint distribution of choice and RT distribution. More generally, the across-trial RL process could be swapped with other dynamic cognitive processes such as Bayesian updating, latent state models of attentional processes or modulatory models of cognitive control. Similarly, the within-trial DDM process can be replaced with any suitable decision models (e.g., Angle DDM, race models such as LBA to account for *n*>2 choice behaviors, etc). Likelihoods for the entire joint RL-SSM process can be differentiable and used with either analytic or likelihood-free methods (e.g., LANs).

#### Box 4.

Centered vs. non-centered parameterizations of hierarchical models

Hierarchical models are most intuitively written in their *centered* form. For a random-intercept regression in which each participant *j* has an intercept *β*_*j*_ drawn around a group mean *μ* with group standard deviation *σ*:

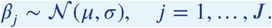

This is conceptually clean but can produce posteriors with awkward geometry: as we explore the posterior space, when we propose small *σ*’s, the conditional distribution over *β*_*j*_ collapses to a point at *μ*, producing a so-called *funnel*. Hamiltonian Monte Carlo samplers — including NUTS (***Hoffman et al., 2014***), the default in PyMC (***Abril-Pla et al., 2023***) — use a single global step size and cannot efficiently navigate both the wide upper region and the narrow neck of this funnel simultaneously, biasing exploration near the apex (***Betancourt and Girolami, 2015***). The problem can be subtle: divergence counts and 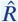 values are sometimes flat enough to look healthy even when the sampler has missed important posterior mass, or you see a high divergence count, even though the chains look ostensibly healthy in turn.

The *non-centered* reparameterization (***Papaspiliopoulos et al., 2007; Gelman, 2004***) samples a standardized offset and applies the location/scale transform after the fact:

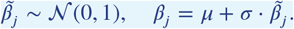

The key is that the two forms describe identical posteriors over *β*_*j*_, but the sampler now explores the standardized 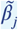, whose geometry is independent of the setting of *σ* — the funnel disappears (or in other words, is undone).

The choice is however not unconditional. Centered tends to be preferable when each group is well informed by the data (the pull of the *β*_*j*_’s toward 0, via small *σ* settings is resisted), while non-centered tends to win when groups are sparsely observed and prior geometry dominates (***Betancourt and Girolami, 2015; Papaspiliopoulos et al., 2007***). `HSSM` inherits Bambi’s default of non-centered parameterization for normal hyper-priors on grouped parameters (***Capretto et al., 2022***) and exposes the choice via the `noncentered` argument.

`HSSM` includes the `RLSSM` class to allow flexible construction of such RLSSM models in a user-friendly manner. Like the SSMs, the class also supports regression models to allow neural or other covariates to inform any model parameter, including the learning process or decision process.

More generally, `HSSM` provides ready abstractions to work with a broad class of models, beyond RLSSMs, that involve longer-timescale, across-trial cognitive processes informing the dynamics of a shorter-timescale, within-trial decision process. This can include Bayesian updating, hidden markov models of task states, optimization of speed accuracy tradeoffs with experience, etc. The analytical or LAN-based likelihoods for the decision process can be flexibly combined with cognitive processes that operate across trials and longitudinally affect the decision process parameters. These can be specified via simple Python callables. All other core `HSSM` features — hierarchical modeling, regression links to external covariates, variational inference, etc. — are naturally inherited.

**Listing 5.**
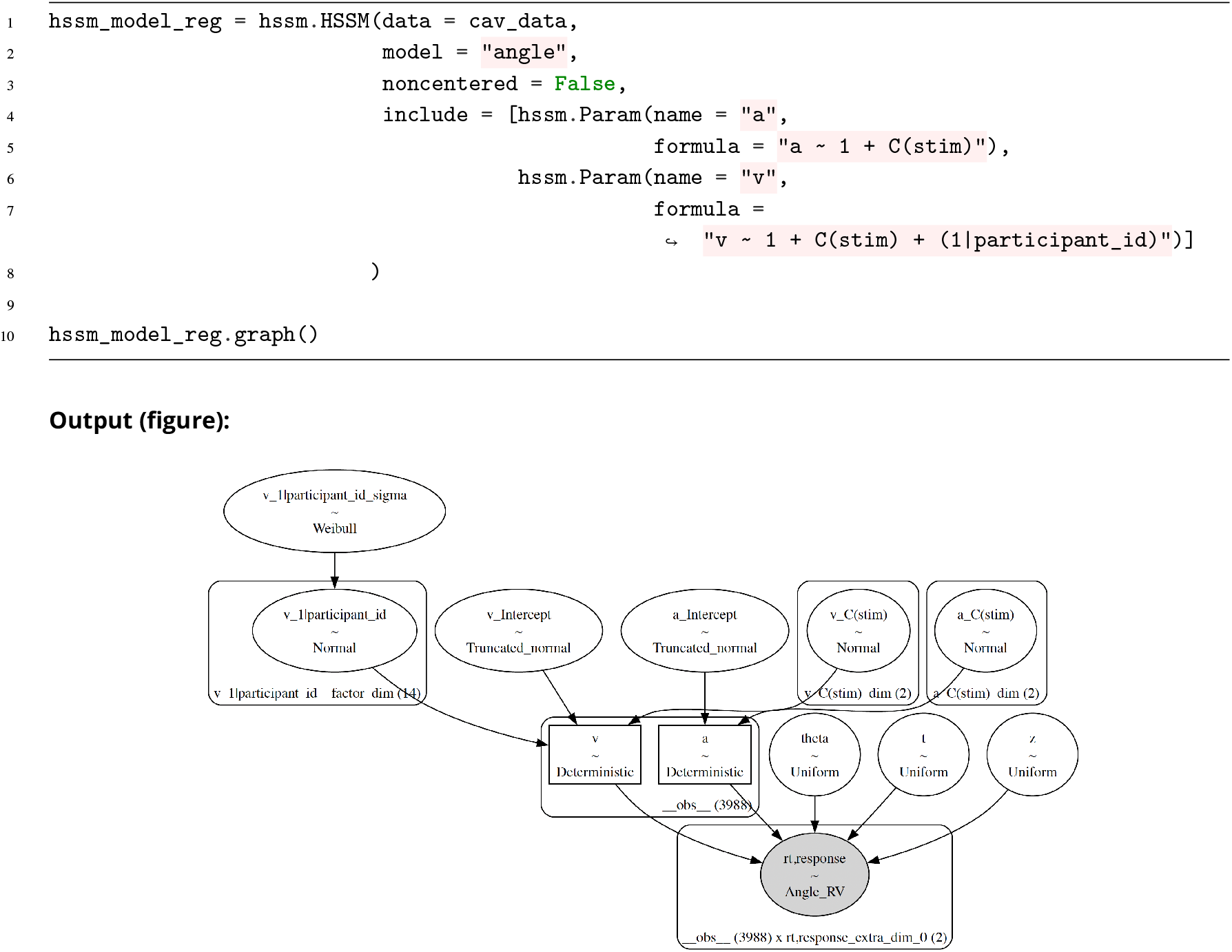
Specifying a hierarchical model with regression backends on the “`a`” and “`v`” parameters of the “`angle`” DDM with `HSSM`. `HSSM` allows multiple approaches to specifying such regressions. Here we show the recommended procedure, via the `include` argument and a list of `hssm.Param` objects.

#### Example

For simple illustration, we consider a two-armed bandit task where, on each trial, a participant selects one of two options, each delivering a binary reward. The task used three reward-contrast conditions, similar to the probabilistic selection task (***Frank et al., 2004***): AB was the easiest condition with 0.90/0.10 reward-probability split between the choices, CD had an intermediate 0.75/0.25 split, and EF was the hardest condition with a weaker 0.60/0.40 split. These probabilities have to be learned from experience with each choice. For each condition, the agent independently maintains value estimates *Q*(*a*_*i*_) for each action, updated after each trial via the delta rule:

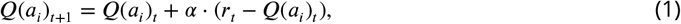

where *α* is the learning rate and *r*_*t*_ is the continuous reward received. Although `HSSM` can support basic RL with softmax choice, here we showcase an example whereby the choice rule has dynamics described by the angle DDM (see Box 3). A single accumulator drifts with Gaussian noise toward one of two absorbing boundaries, with the drift rate on trial *t* set proportional to the current value difference:

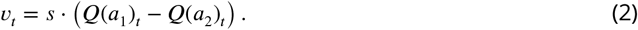

The user specifies a learning process function that computes trial-wise decision parameters from the model’s free parameters and the observed trial history. `HSSM` then evaluates the decision process log-likelihood (whether via surrogate likelihood or analytical form) over the resulting parameter trajectories. The input `DataFrame` must contain columns for response time (`rt`), response label (`response`), and any additional fields required by the learning process (e.g., `feedback`). A participant identifier column (`participant_id`) is also required. The trials should be ordered correctly to reflect the observed sequence of stimuli and feedback presentation. Listing 6 shows how this model is constructed. Once the RLSSM model is constructed, fitting proceeds via the standard `.sample()` interface inherited from the base `HSSM` class, as shown in Listing 7. Figure 5 shows how RLSSM jointly captures the choice and RT distributions in the dataset.

**Figure 5.**
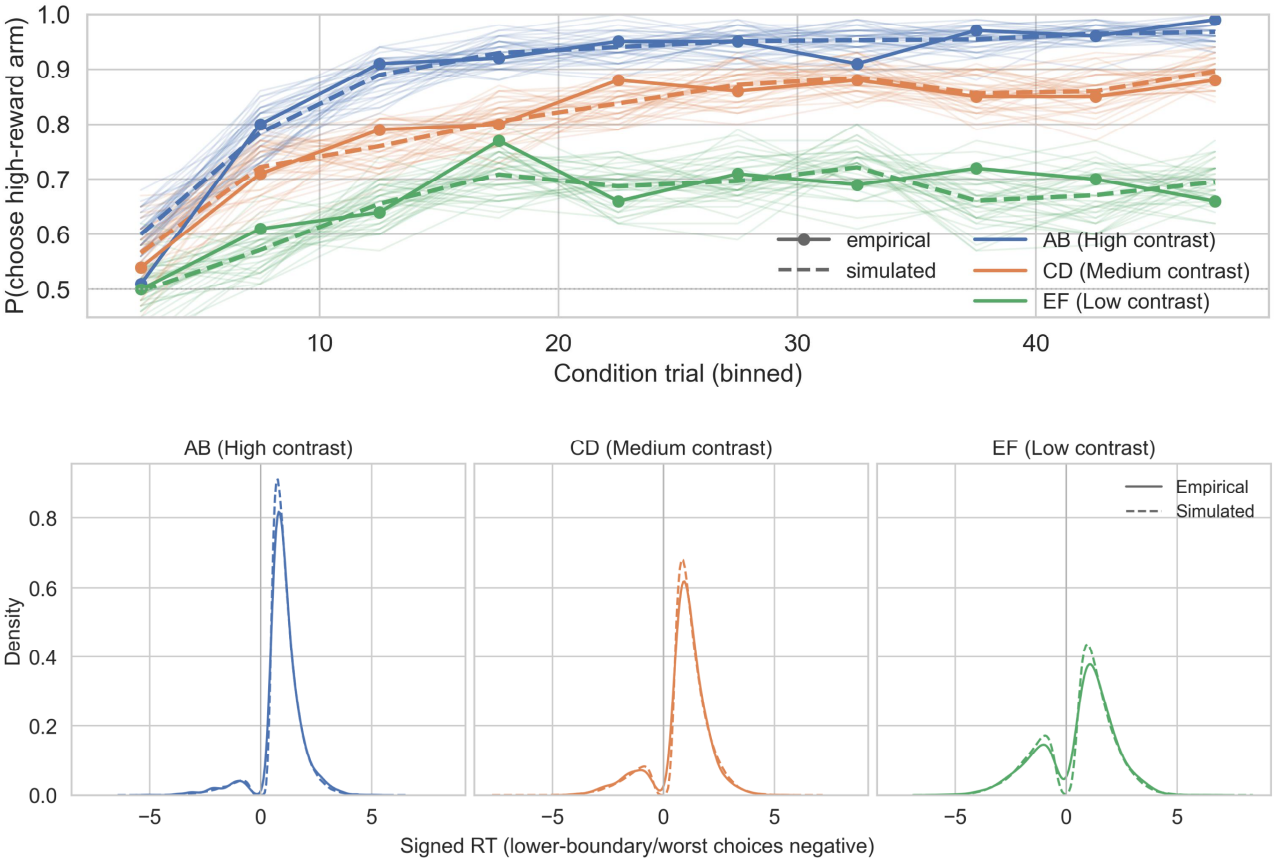
Posterior predictive checks for the RLSSM model. Here the Q learning rule is used and the decision process is Angle (linear collapsing bound DDM), but either process can be replaced with other learning rules or SSMs. Top: early-trial choice accuracy, shown as the probability of choosing the high-reward option across binned condition trials. Empirical trajectories are shown with solid lines and markers, while posterior predictive simulations are shown with dashed lines; thin translucent lines show individual simulated datasets. Bottom: condition-wise signed response-time distributions, where upper-boundary/best-option choices are positive and lower-boundary/worst-reward choices are negative. Across AB, CD, and EF reward-contrast conditions, the model captures both the learning-related separation in choice accuracy and the broad structure of the response-time distributions.

**Listing 6.**
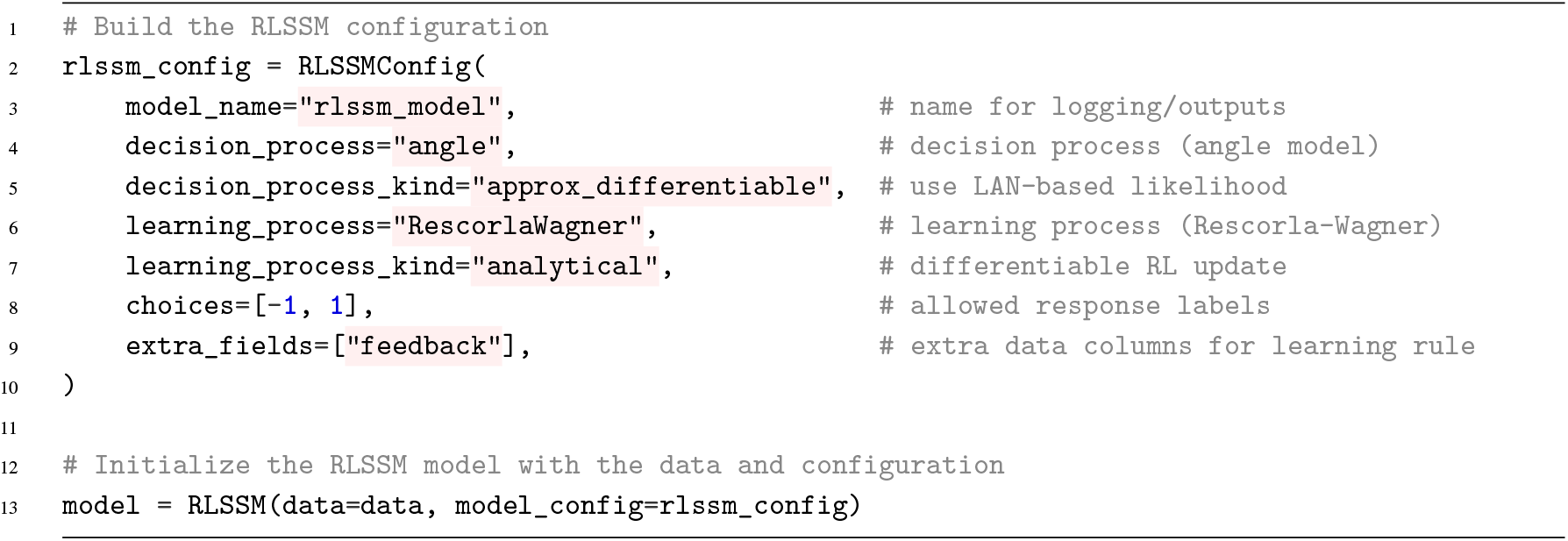
Basic `RLSSM` model configuration in `HSSM`. The `RLSSMConfig` object ties together the choice of decision process (here, the angle DDM with an approximate differentiable likelihood), the learning process (here, delta rule with an analytical update), and the data columns required by the learning rule.

**Listing 7.**
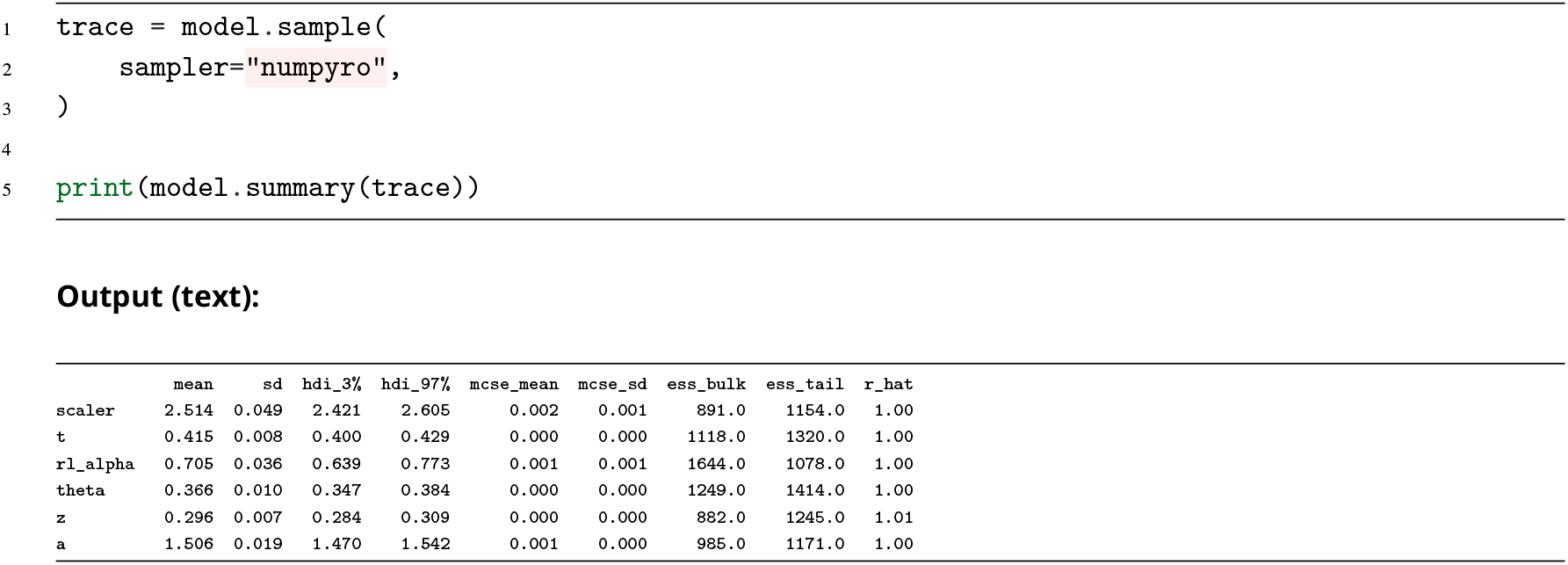
Fitting an RLSSM model with the `numpyro` backend, and inspecting the posterior summary. The hierarchical regression interface, NUTS samplers, and ArviZ-compatible outputs are all inherited unchanged from the core `HSSM` class.

### Native Plots

Beyond the ability to interface directly with the large arsenal of diagnostic plots that ArviZ (***Kumar et al., 2019***) provides, `HSSM` includes three types of plots to help with sequential sampling models.

The first is the basic `.plot_predictive` function. The plot is freely configurable and allows us to plot the basic choice-RT distributions from n-choice models. If we are dealing with a two choice model, RTs for which the choice was −1 (i.e. lower boundary choices), will be plotted on the negative RT axis. For models which more than two choices, color determines choice and all distributions are on the positive interval. The plot can naturally be used for both plotting the prior as well as the posterior predictive. By default it uses as parameters the mean prior/posterior, however it allows explicit representation of uncertainty.

**Listing 8.**
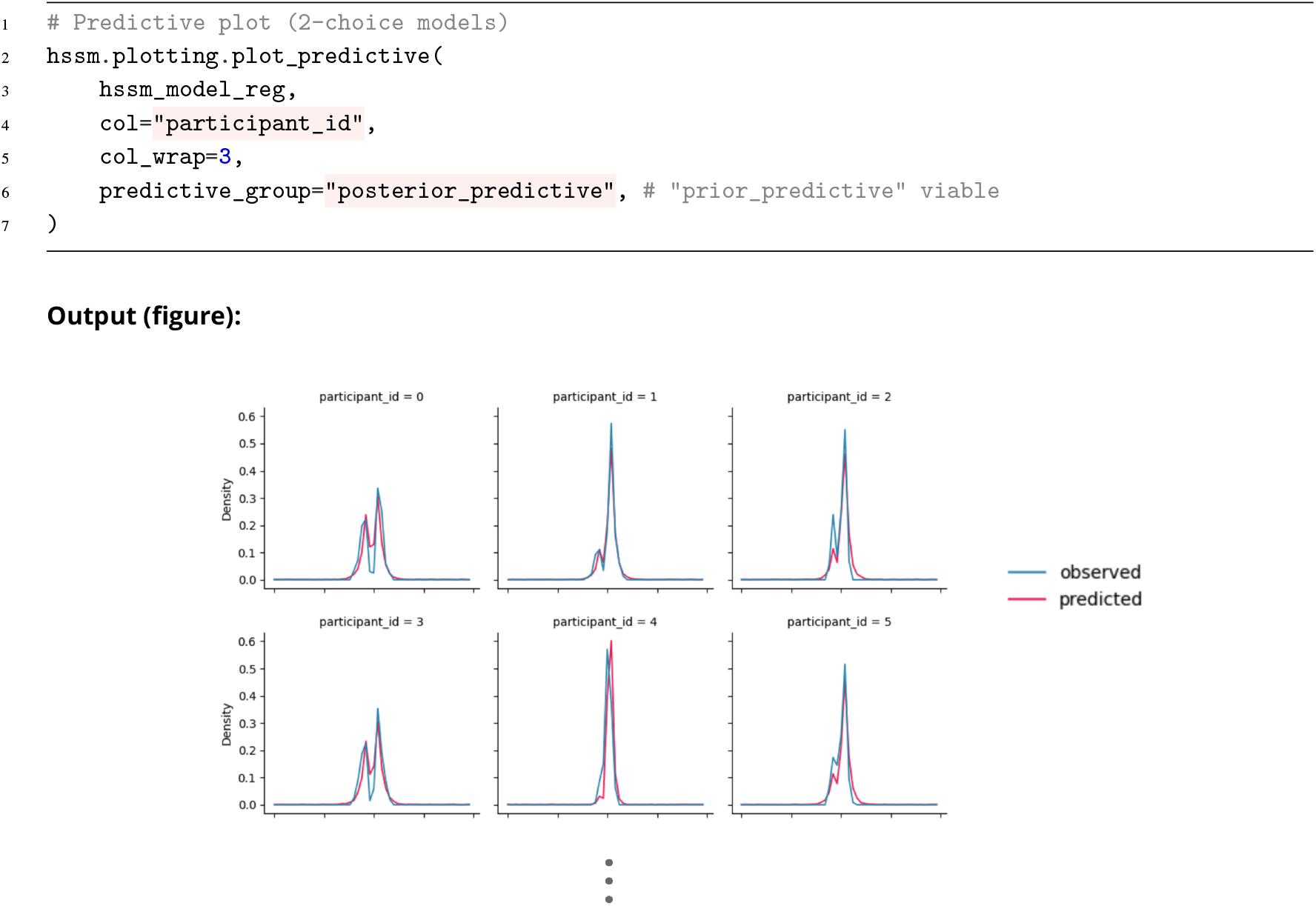
Basic posterior predictive plots for 2-choice models. The `plot_predictive` method allows for large stylistic and computational flexibility. We highlight the `predictive_group` argument, which specifies whether we would like plot the prior or the posterior predictive.

Second, `HSSM` ships with a configurable quantile probability plot (***Ratcliff and Tuerlinckx, 2002***), via the `.plot_quantile_probability()` method. Quantile probability plots are designed to show how choices and RT distributions vary by task condition (see `WL`, `LL`, `WW` on top of the x-axis in Listing 9, rt quantiles. Quantile probability plots are widely applied diagnostic tools in the context of sequential sampling models and help parcel out, with a single glance, the joint changes in reaction time distributions and choice probabilities, when moving across different conditions in a given experiment.

For each condition we have two choices with respective choice proportions (`Proportion` on the a-axis). For each choice we then have the quantiles for the corresponding reaction time distribution. The dotted lines here represent the real data. (In the case of continuous experimental covariates one can still display these into “conditions” here, by simply binning the data into n-levels.)

The `plot_quantile_probability()` method then allows us to plot whether these patterns of choices and RT distributions are expected from the model given the uncertainty over its parameters. We can pick the type of visualization (“`ellipse`” was chosen in our illustration), where the ellipse shows the 95% highest density interval on where the data are expected to lie (again both choice proportions and corresponding RT quantiles) for each task condition. The model is a good fit if the data lie within the ellipses. While we preferably want the ellipses to form a tight bound around their corresponding data-points, many factors, such as the number of trials per participant, the group level parameter variance across participants etc., play a role in what an ‘expected good outcome’ may be in a particular modeling situation.

Listing 9 shows an example, based on our `cav_data` dataset. Figure 6 shows an example that contrasts two model fits, where the underlying data is simulated with collapsing bounds, and two separate models are then fit to the data. Ignoring the collapsing boundaries in this example shifts the mass of the incorrect choice distributions away from the data distribution. This is because the fitted fixed boundary has to adjust to compensate for the true collapsing boundary, and thus errors are expected to be made in earlier RT quantiles than are consistent with the data. Thus, QP plots are useful for visualizing how data may be expected under posterior uncertainty, but they are also helpful for identifying where model fits may be lacking.

**Figure 6.**
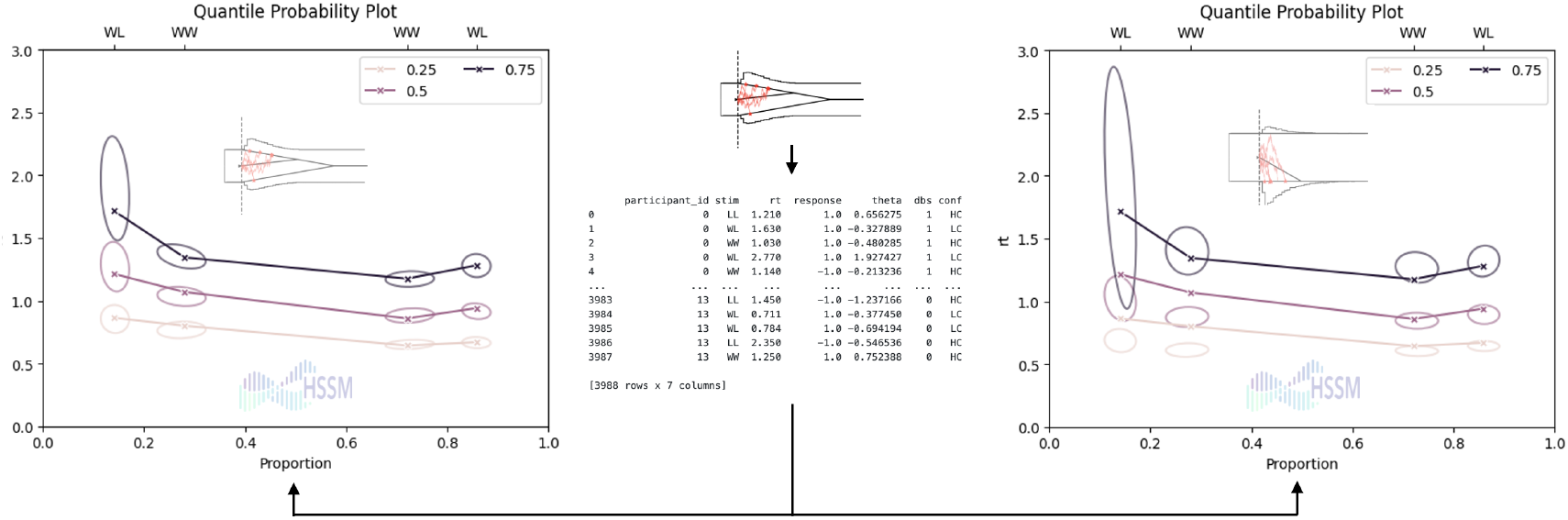
Quantile probability plots for visualizing model fits to data, comparing a true and misspecified model. This figure illustrates how QP plots can identify features of the data that the model may predict or miss. Here data is generated synthetically from a model with linearly collapsing boundaries, and then fit with that same model (left) or with the simple DDM (right). The ellipses, reflect the uncertainty (HDI) of where the data are expected to lie for both choice probability in each condition (x-axis) and corresponding RT quantiles (y-axis). For the true model, the data are well within the expected uncertainty of the model for both choice and RT quantiles in each condition. For the misspecified model (right), the model misses some of the RT quantiles, in particlar error are expected to be faster than observed empirically. While synthetic, to maintain harmony with the rest of our examples, the data here is otherwise structured as the “`cavanagh_theta`” dataset.

**Listing 9.**
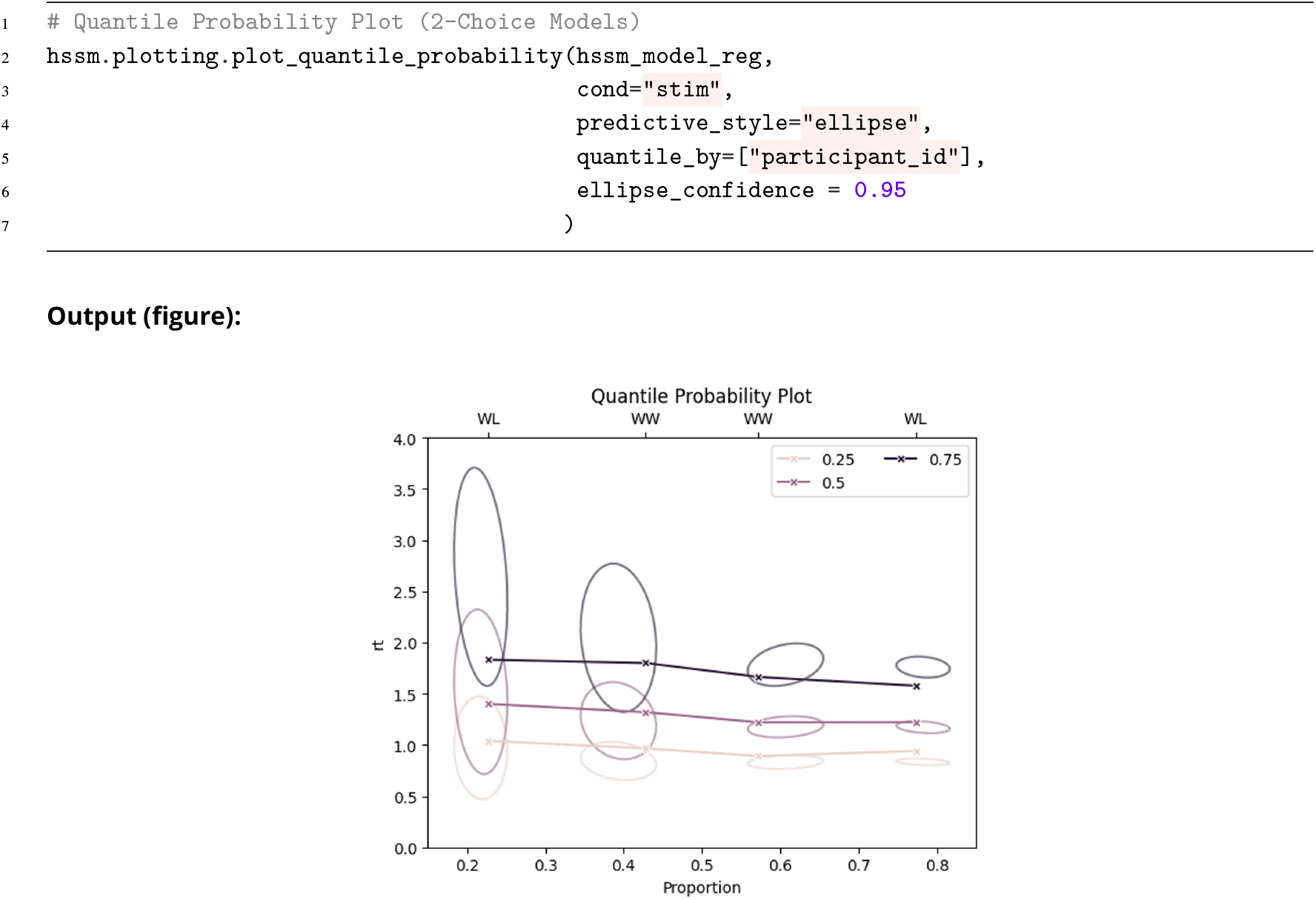
`HSSM` ships with quantile probability plots (***Ratcliff and Tuerlinckx, 2002***), via the `plot_quantile_probability` method. The plot has various computational and visual knobs that can be adjusted to specialize it to a particular scenario. We highlight the get `quantile_by` argument, which establishes at which level quantiles are being computed. If e.g. (as illustrated) the argument is set to “`participant_id`”, we compute quantiles first per participant, then aggregate. The ellipses reflect the uncertainty of the quantiles and choice probabilities in line with the chosen aggregation process.

Third, `HSSM` comes with, what we call, *model cartoon plots* via the `plot_model_cartoon()` function. These plots integrate a visual of the underlying cognitive process model with the posterior predictive. As the other two plots, model cartoon plots are highly customizable. Users can turn on and off various elements of the cognitive process model, decide whether or not to include posterior uncertainty as a plot element. Listing 10 provides an example using the `cav_data` dataset. Figure 7 provides a didactic overview of the construction of the model cartoon plots, to aid comprehensive at a glance.

**Figure 7.**
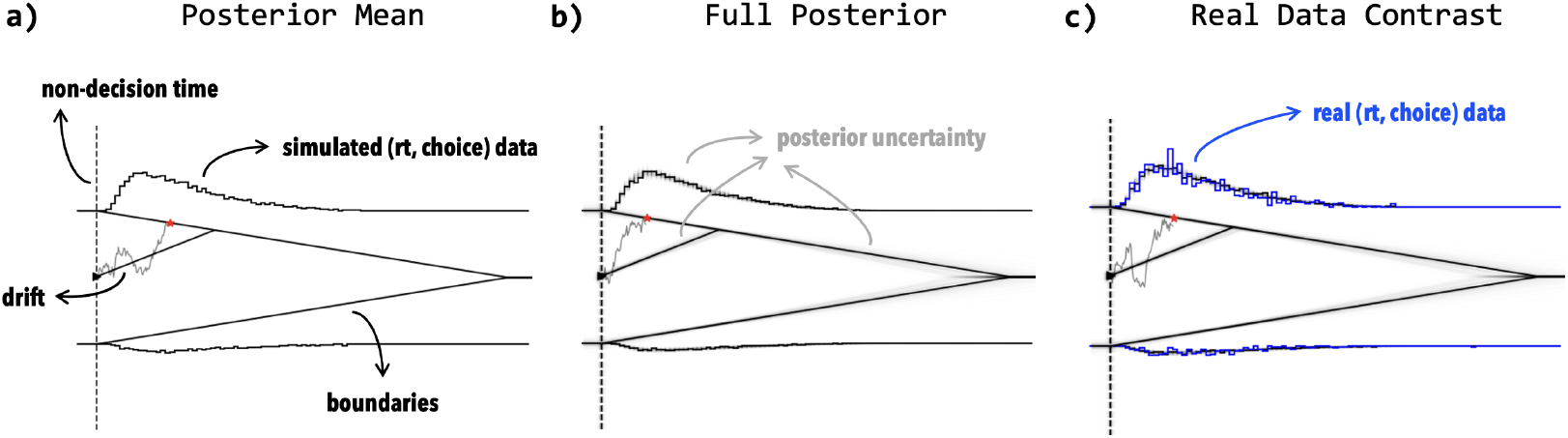
Anatomy of the model cartoon plot. The plot is built in layers. **(a)** The structural elements of the cognitive process model are overlaid on the response time axis: here, the decision boundaries, non-decision time (dashed line), drift trajectory (with boundary crossing marked by a red star), and the choice-conditioned response time histogram evaluated at the posterior mean. **(b)** Posterior uncertainty is added by overlaying many samples from the posterior predictive distribution (grey shading). **(c)** The observed data are added in blue, allowing direct visual comparison with the posterior predictive. Each element can be toggled independently via keyword arguments.

### Advanced: Low-level API

We include in `HSSM` a low level interface for advanced users, who may wish to work directly with PyMC. This includes two types of utilities. First, pre-constructed random variables, for named models shipped with the toolbox. These can then naively be used in a given unrestricted PyMC model. This is illustrated in the figure and code in Listing 11. Second, we provide convenience functions so that users can, with few lines of code, construct their own PyMC random variable, by putting together a simulator (optional, but convenient to let PyMC automatically handle prior and posterior predictive sampling) and a likelihood function. These utilities are designed to be easy to use especially with simulators from the `ssm-simulators` package and likelihoods derived from `LANFactory`, however they are not restricted to this setting. We show multiple examples for each case in the documentation. The likelihood constructors, can themselves downstream be used for fully custom workflows and fitting procedures.

### Advanced: Construct Model and Exit the Ecosystem

As expressed earlier, we treat `HSSM` as a modular component in scientific workflows. If desired a very large fraction of common workflows for computational cognitive process modeling can be covered via the `HSSM` ecosystem. However, if a user finds a particular component useful, we try to emphasize entry and exit points so that the component can be used in isolation, instead of attempting to lock users into more aspects of our vision that they wish to partake in.

**Listing 10.**
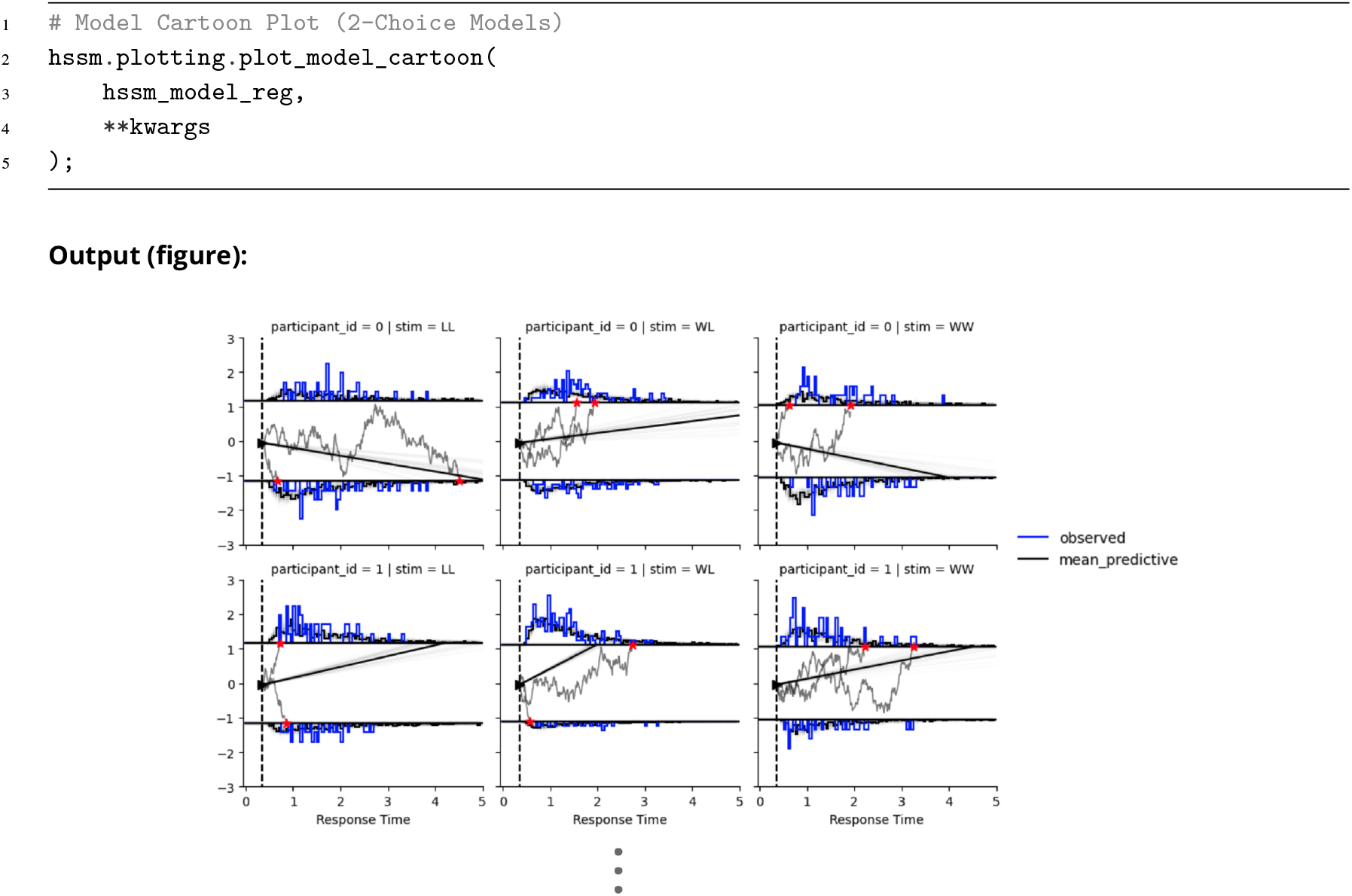
Model cartoon plot for a two-choice SSM. The `plot_model_cartoon()` function visualizes the fitted cognitive process model alongside the observed data. Blue histograms show observed response time distributions (plotted on the negative axis when the choice was −1 and on the positive axis otherwise). Overlaid in grey are individual sampled decision trajectories from the fitted model, with boundary crossings marked in red. The black curve traces the mean posterior predictive distribution. Dashed vertical lines indicate the non-decision time. Users can toggle trajectories, posterior uncertainty, axis layout and various other stylistic elements via the keyword arguments.

**Listing 11.**
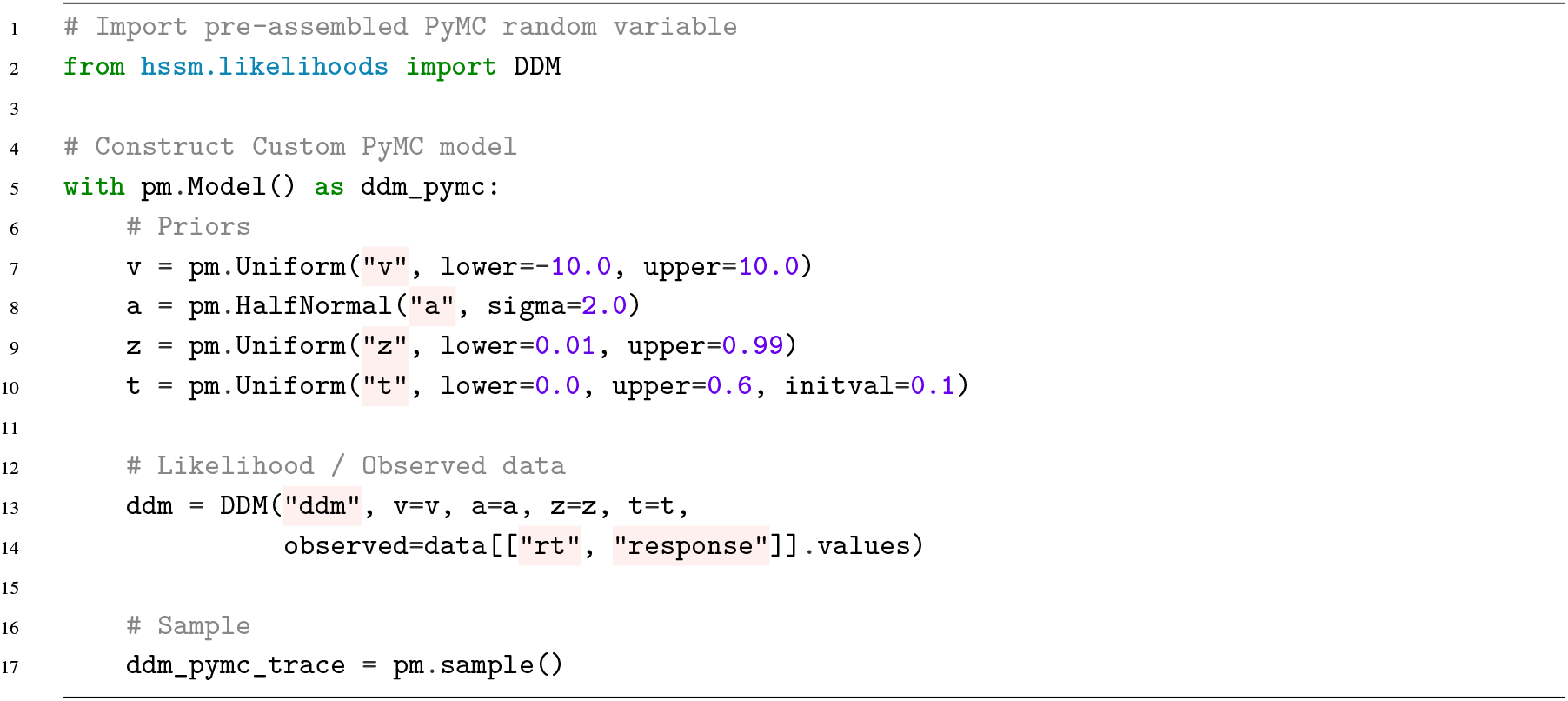
Low level API. This listing shows how we can use random variables that are shipped with HSSM directly inside custom PyMC models. Advanced users can use this lower level interface to build much more ambitious models than feasible via the current base `HSSM` class.

**Listing 12.**
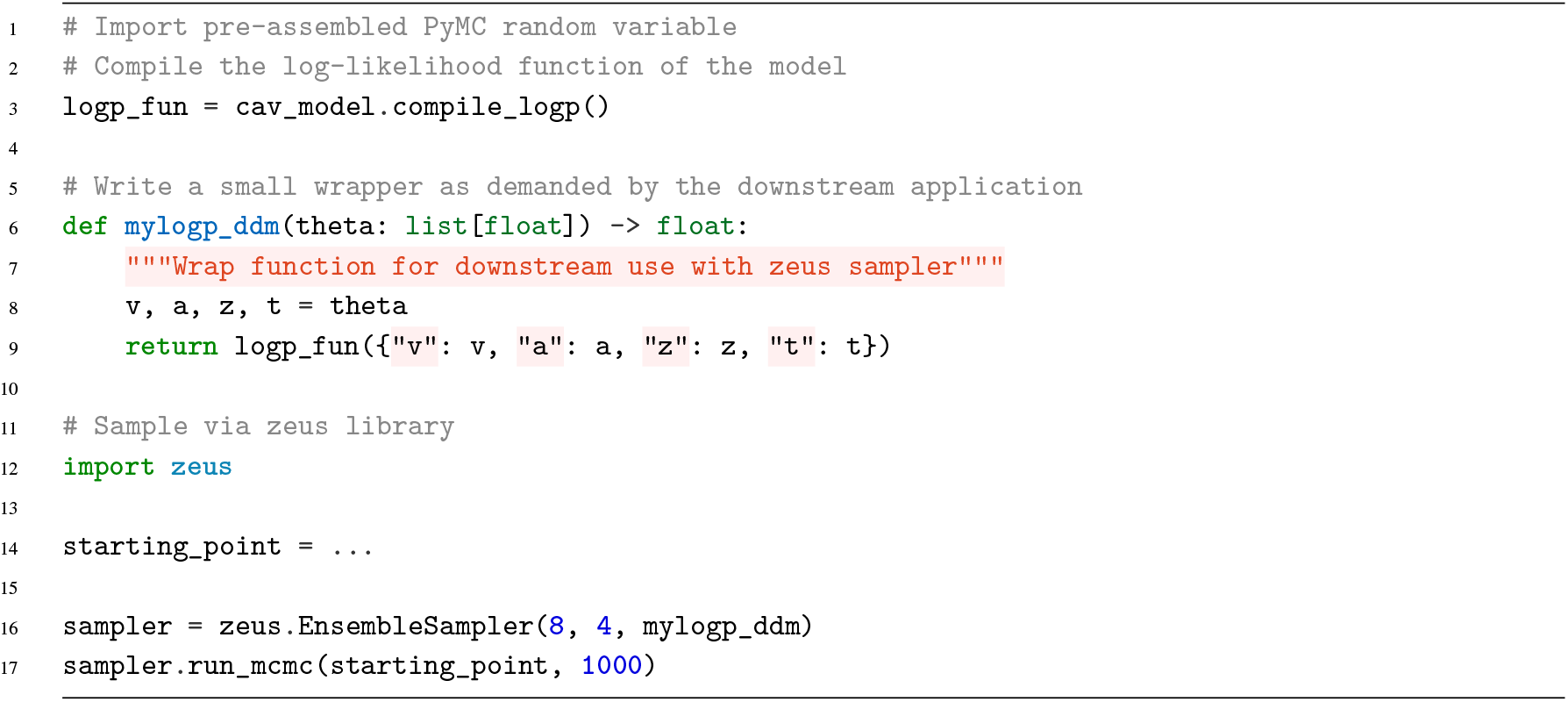
External samplers. This listing showcase how we can compile the log-likelihood function of a given `HSSM` model (we are accessing the underlying PyMC model directly) and downstream use it to serve as the basic likelihood to be called through entirely different MCMC sampler libraries. The `zeus` library serves as a convenient example to showcase how we can seamlessly exit the entire PyMC ecosystem while making `HSSM` do useful groundwork for us nevertheless.

The code example above illustrates how to use `HSSM` as a model construction engine, however then leave the framework to use e.g. a completely different sampling engine downstream, which may even live entirely out of the broader PyMC ecosystem. The key is that we can exploit `HSSM` to construct the underlying model, compile the graph as a function and simply use this function to our liking downstream. We show an example, in which we interface with the `zeus` library for MCMC sampling (***Karamanis et al., 2021***) after model construction.

## ssm-simulators

### Place in the ecosystem

Within the `HSSM` ecosystem, `ssm-simulators` serves as the foundational layer upon which the other packages are built. Its simulators provide the forward models that `HSSM`uses for prior and posterior predictive sampling, and its training data generation pipelines produce the structured datasets that `LANFactory` consumes to train surrogate likelihood networks.

Beyond this technical role, `ssm-simulators` is the natural entry point for community contributions to the ecosystem. Theoreticians who wish to make a new cognitive process model available to the broader research community can implement it here. The path to downstream availability is then streamlined: if analytic likelihoods are not available, training data can be generated via the built-in pipeline, a likelihood surrogate can be trained via `LANFactory`, and the resulting model becomes immediately testable against experimental data via `HSSM`.

We note that `ssm-simulators` is equally usable as a standalone simulation library. Researchers can simply use it to explore model behavior and generate synthetic datasets to be used downstream for any task, without needing to engage with any other aspect of the `HSSM` ecosystem.

### Core Functions

Following the mermaid diagram in Figure 8, the `ssm-simulators` package is meant to achieve the following two core functionalities. First, this package serves as the basic simulation engine we use in the `HSSM` ecosystem. The core simulators are accessible via the `Simulator` class.

**Figure 8.**
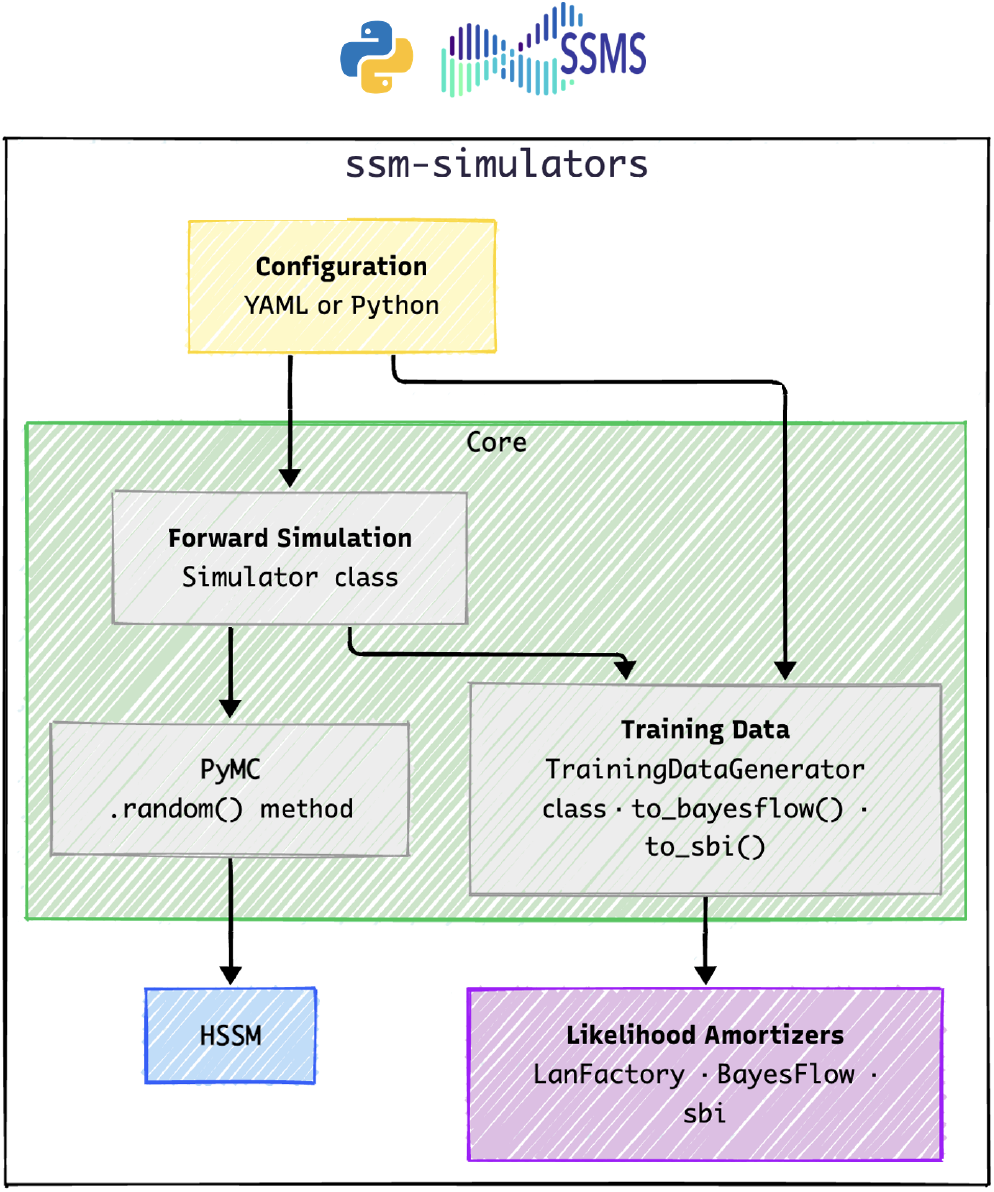
Architecture of the ssm-simulators package. The package provides two core functionalities: forward simulation of cognitive process models via the `Simulator` class, and a modular training data generation pipeline that combines configurable likelihood estimators and sampling strategies to produce structured datasets for downstream likelihood approximation (e.g. via `LANFactory`, connectors to the `sbi` (***Tejero-Cantero et al., 2020***) and the `BayesFlow` (***Radev et al., 2020***) libraries exist). **Figure 8—figure supplement 1**. `ssm-simulators` Conceptual Architecture: Detailed

Via this class we are able to access a wide range of pre-implemented (and extensible list of) cognitive process models. Below is a basic example:

The second core function of this library is to provide a *training data generation pipeline*, to produce structured datasets that can be used for visualization or model behavior and, when needed, Neural Network training for purposes of likelihood approximation. This functionality is implemented in a modular manner. The user passes configurable components (*Estimators* and *Training Data Strategies*) into a training data pipeline, represented by the `TrainingDataGenerator` class. *Estimators* here represent an approach to get training signals for likelihood evaluations: `ssm-simulators` includes two types of *Estimators* out of the box. The respective abstractions invite the community to provide others. A *Training Data Strategy* is an algorithm to sample (*rt, c*) pairs for evaluation by the `Estimator`.

To provide a concrete example of this distinction, an `Estimator` is an algorithm to provide likelihood evaluations that can be used at training time. For a given parameter set, we might construct a KDE (***Węglarczyk, 2018***) from simulations, as proposed for the original LAN training pipelines (***Fengler et al., 2021***). This KDE can then be evaluated for (*rt, c*) pairs, which opens up a rather large set of options. Should we sample from the process itself? Should we evenly distribute over choices *c* and evaluate *rt* with uniform spacing in some interval [0, *rt*_*max*_)? The *Training Data Strategy* provides a concrete decision on how to query a constructed `Estimator` on a given parameter set. Together, the `Estimator` and the *Training Data Strategy* provide a complete training data pipeline for which the `TrainingDataGenerator` acts as an orchestrator. Below is a simple example to invoke each of the core functionalities. Figure 8 provides a conceptual breakdown of the package design.

**Listing 13.**
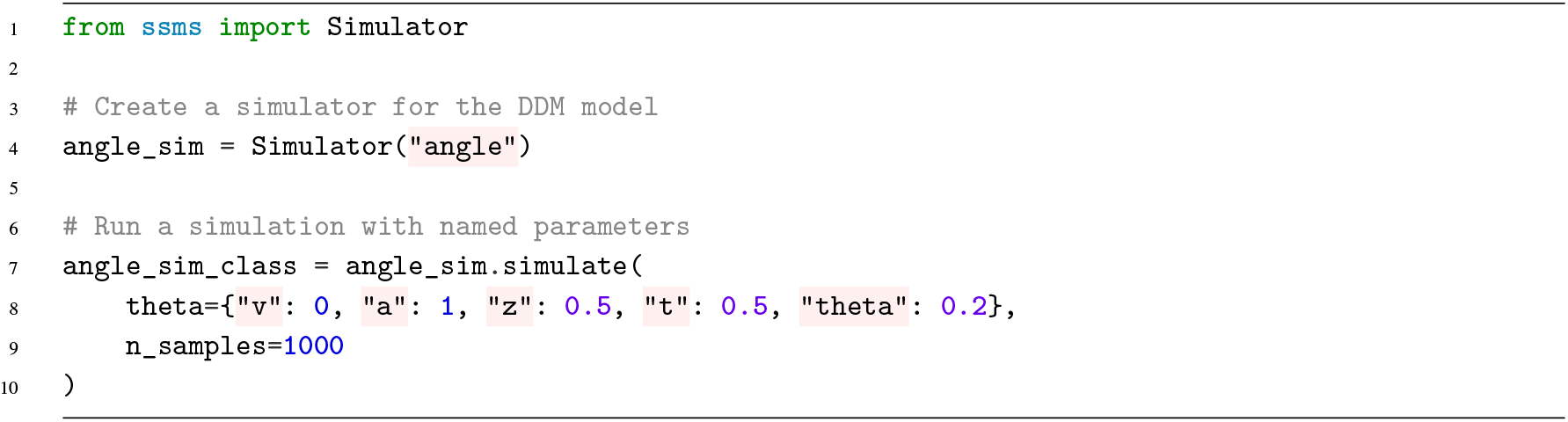
Simulation of the angle DDM using `ssm-simulators`

Listing 13 shows how to run a basic simulation. The logic is simple. First, the user instantiates a `Simulator` class, passing the name of a registered model. The user can then invoke the `.simulate()` classmethod to perform simulations. The `theta` argument, which allows the passing of model parameters, is flexible and allows users to pass scalar or vectors for each parameter, via a `numpy arrays` or a dictionary with parameter names as keys. If the user passes a vector for any parameter, the simulator treats the length of this vector as the number of *trials* and `n_samples` will act as the number of samples for each trial. The `sim_out` variable is a Python dictionary with three keys: “`rts`” and “`choices`” which hold the core simulation output and “`metadata`” which holds a host of extra information about the simulator call itself to help with identification downstream.

### Example: Training Data Generation

**Listing 14.**
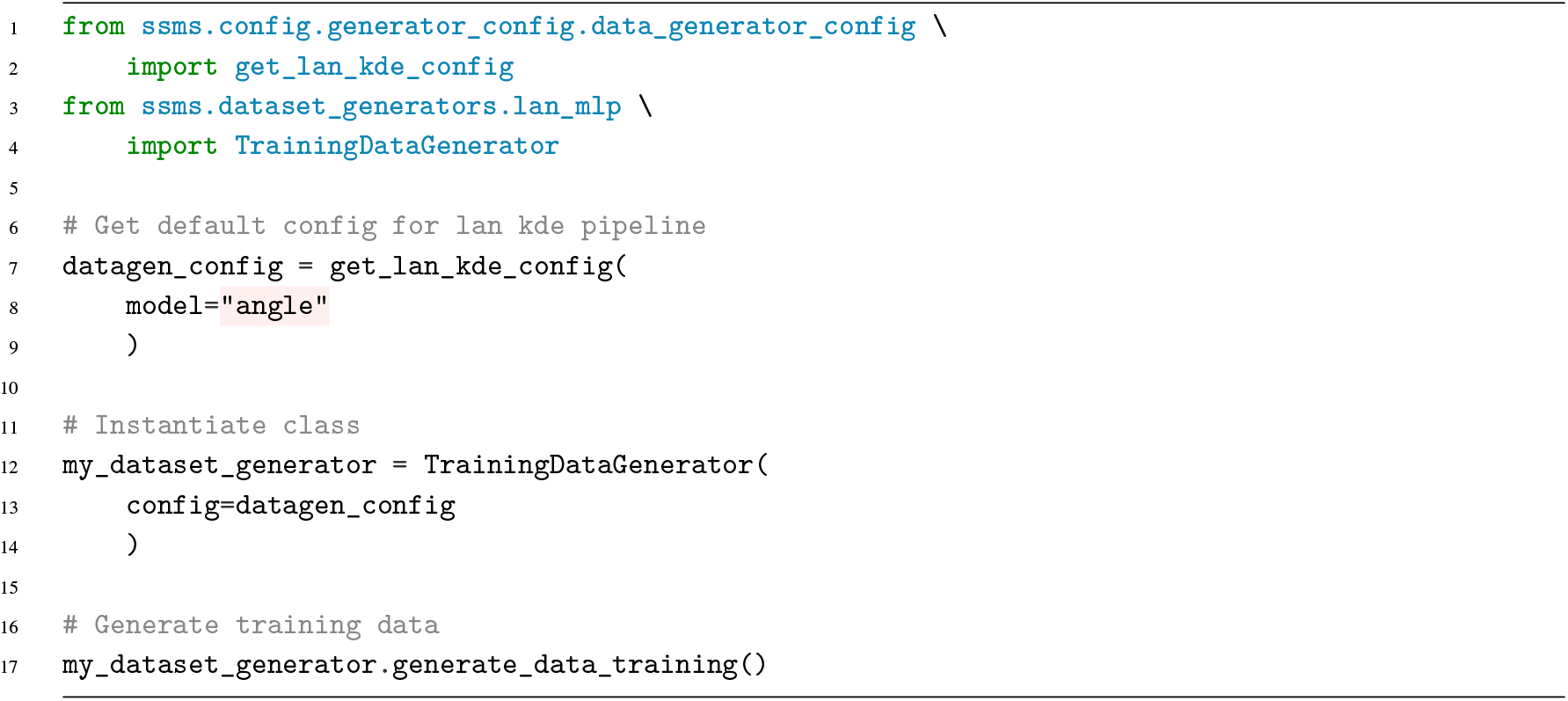
Basic training data generation for the “angle” DDM with `ssm-simulators`

Listing 14 provides a simple way to run training data generation for a basic default pipeline, which uses KDE based estimators to generate training labels.

### Example: Training Data Generation via CLI

The `ssm-simulator` package comes with a simple YAML configurable command line interface, which helps users create training data from registered models with a highly simplified interface. Listing 15 shows the simple terminal command to launch training data generation.

**Listing 15.**
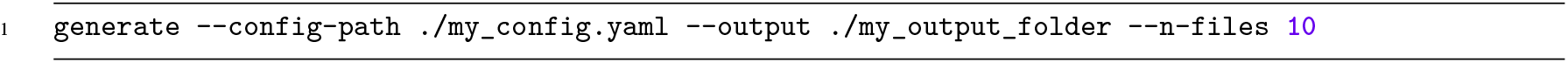
`ssm-simulators` CLI interface for training data generation from YAML configuration files.

## LANFactory

### Place in the Ecosystem

`LANFactory` occupies the middle layer of the ecosystem, bridging the gap between simulation and inference. It consumes training data produced by `ssm-simulators` and outputs ONNX-formatted (***Bai et al., 2019***) neural networks that`HSSM` can download and deploy as differentiable surrogate likelihoods. The standardized pipeline from training through ONNX conversion to HuggingFace upload ensures that any new model contributed via `ssm-simulators` can be made inference-ready with minimal friction. As with the other ecosystem components, `LANFactory` is designed to be optional rather than obligatory. Users who obtain likelihood approximations through other means — whether from analytical derivations, numerical solvers such as `PyDDM` (***Shinn et al., 2020***), or alternative neural density estimators such as those from `BayesFlow` (***Radev et al., 2020***) — can bypass LANFactory entirely and supply their likelihoods directly to `HSSM`. In this sense, `LANFactory` provides the default and most streamlined path for contributing surrogate likelihoods, but it is not a gatekeeper.

### Core Functionality

The `LANFactory` package provides the glue between the the training data generators in `ssm-simulators` and the likelihood constructors in `HSSM`.

The package is responsible for training neural networks that serve as surrogate likelihood evaluators for analytically intractable cognitive process models. It accepts structured training data — typically produced by the `ssm-simulators` package — in which parameter configurations are paired with estimated log-likelihood values at sampled (rt,c) points, and uses these to train lightweight feedforward networks via either JAX or PyTorch.

The package supports three network types: likelihood approximation networks (LANs), which approximate the full log-likelihood surface; choice probability networks (CPNs), which estimate the probability of each choice given model parameters; and omission probability networks (OPNs), a special case of CPNs used for modeling omission trials (i.e., when no response is recorded) (***Leng et al., 2024***). Once trained, networks are converted to the ONNX format for framework-agnostic deployment and can be uploaded to the ecosystem’s HuggingFace repository, from which`HSSM` retrieves them automatically at model construction time.

In keeping with the ecosystem’s modular design philosophy, `LANFactory` is built to be extensible. Users can incorporate custom network architectures beyond the default feedforward design, and the training loop is configurable via YAML files or direct Python API calls. The package is not, however, a prerequisite for contributing likelihoods to `HSSM` — it provides the most streamlined path for doing so, while leaving room for alternative approaches. Figure 9 provides a conceptual breakdown of the package design.

**Figure 9.**
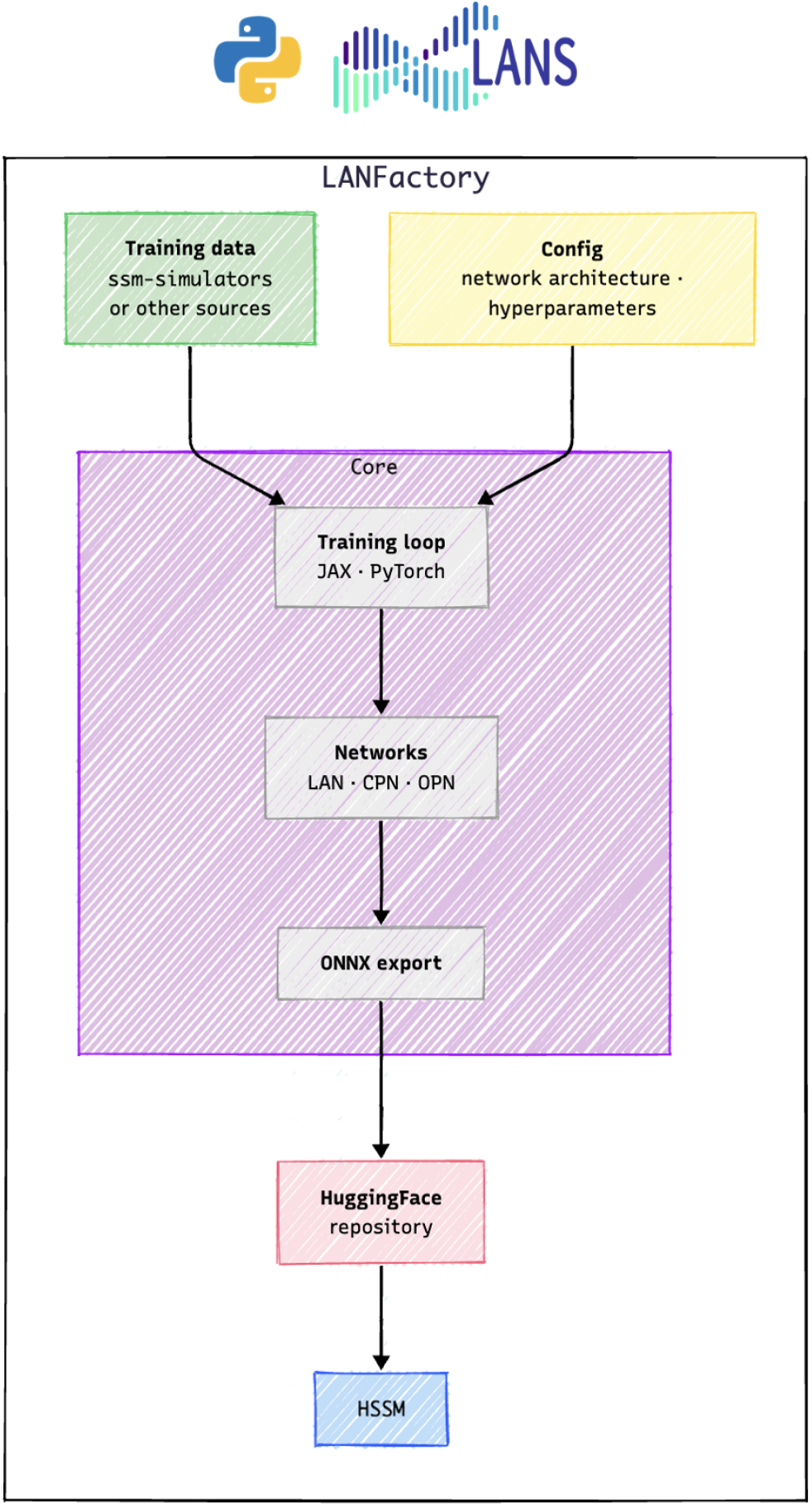
Architecture of the `LANFactory` package. The package trains surrogate likelihood networks (LANs, CPNs, OPNs) from training data produced by `ssm-simulators`, using either JAX or PyTorch backends. Trained networks are converted to the ONNX format and uploaded to HuggingFace for deployment within `HSSM` or elsewhere. **Figure 9—figure supplement 1**. `LANFactory` conceptual architecture, detailed.

### Example: CLI Training Workflow

The `LANFactory` documentation includes a full set of tutorials on how to train each type of network via both JAX as well as PyTorch. While the pure python interface remains low code for end-users, to keep code snippets succinct in this illustrative paper, we will only show the CLI workflow, which is exceedingly simple.

We have three steps in the process: First, *train the network* for which we have the `jaxtrain` and `torchtrain` commands. Second, *transform to ONNX* for which we have the `transform-onnx` command. Third, *upload to HuggingFace*, for which we finally have the `upload-huggingface` command.

Network training produces two core files per training run:

1. A *network configuration*
2. A *set of trained weights*

These get passed to `transform-onnx` to produce a single `.onnx` file. The HuggingFace upload finally includes the `.onnx` file, the network**\_**config.pickle file and a *Model Card*, a string description of about the network and underlying simulator with a simple usage example for `HSSM`.

**Listing 16.**
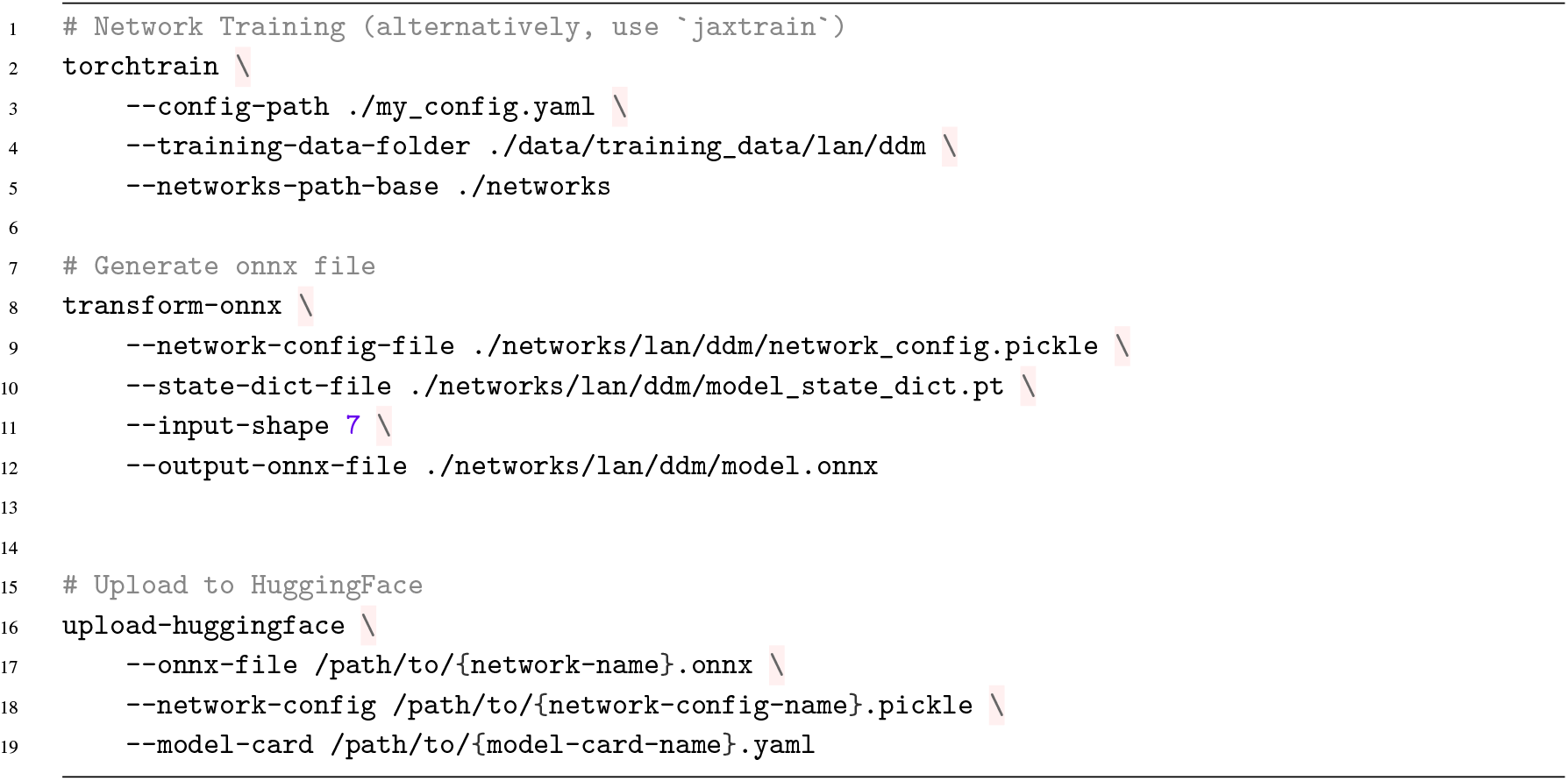
`LANFactory` CLI interface. Three core commands (i) `torchtrain` (or the JAX equivalent) to train the network (ii) `transform-onnx` to generate the ONNX files and (iii) `upload-huggingface` to upload the network to our HuggingFace database.

**Listing 17.**
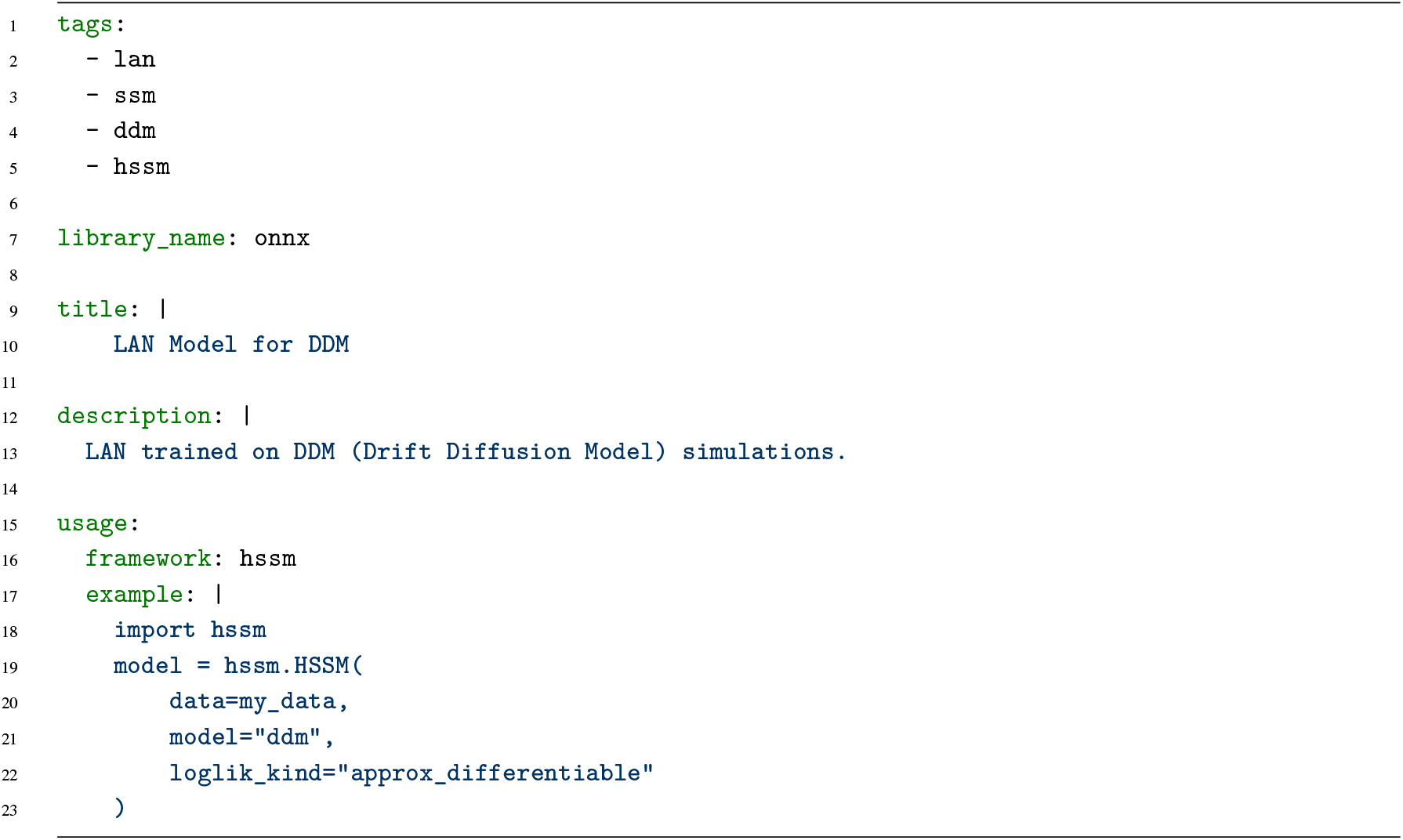
`LANFactory` Model Card. A model card is associated with every uploaded model in the HuggingFace database.

## Related Work

The widespread adoption of sequential sampling models in cognitive neuroscience has been catalyzed by the availability of some accessible software toolboxes (***Wiecki et al., 2014; Ahn et al., 2017; Shinn et al., 2020***). However, as the field has advanced toward larger datasets, more complex hierarchical models, the integration of neural and learning dynamics, and more and more variations of the underlying cognitive process models, a larger strain is being put on the existing package ecosystem to keep up with the demands of experimentalists. We review what we consider contemporarily the most relevant alternatives to `HSSM` below, organized by the methodological niche each occupies, and summarize the comparison in Table 1.

**Table 1.** Feature comparison of likelihood-based software packages for cognitive process modeling. · = fully supported, ◦ = partial or limited support, – = not supported. “Hierarchical Models” refers to full hierarchical estimation with group and subject parameters. “Trial-by-trial covariates” refers to incorporation of continuous trial-level measures (e.g., neural activity, learning signals) as predictors of model parameters. “Full LMER” refers to lmer-style formula-based specification of between- and within-subject effects on model parameters, including random slopes and intercepts. “RL-SSM integration” refers to built-in support for combining reinforcement learning with sequential sampling models. “Surrogate models” refers to support for models lacking closed-form likelihoods via learned likelihood approximations (e.g., neural networks). “User Extensible” refers to the ability for users to add new models or likelihood functions without modifying the package source code. BayesFlow, which targets amortized posterior estimation and surrogate likelihood training via a fundamentally different paradigm, is discussed in the text but omitted from this comparison.

| Feature | HSSM | HDDM | EMC2 | PyDDM | PyBEAM | hBayesDM | rlssm |
| --- | --- | --- | --- | --- | --- | --- | --- |
| Interface language | Python | Python | R | Python | Python | R/Python | Python |
| Bayesian | Yes | Yes | Yes | No | Yes | Yes | Yes |
| Hierarchical Models | • | • | • | – | ◦ | • | • |
| Trial-by-trial covariates | • | • | ◦ | – | – | • | – |
| Full LMER | • | ◦ | • | – | – | – | – |
| RL-SSM integration | • | ◦ | ◦ | – | – | – | • |
| GPU acceleration | • | – | – | – | – | – | – |
| Surrogate models | • | ◦ | ◦ | – | – | – | – |
| User Extensible | • | ◦ | ◦ | • | ◦ | ◦ | – |

### HDDM: The predecessor

For the past decade, the standard for hierarchical Bayesian modeling with SSMs was the `HDDM` (***Wiecki et al., 2014***) Python toolbox, developed in our own lab and used in over 1000 published studies. `HDDM` pioneered hierarchical Bayesian estimation of drift diffusion models in Python, introduced support for trial-by-trial neural regressors down the line, and was later extended with LANs to support a broader class of SSMs and reinforcement learning variants (***Fengler et al., 2022***). However, `HDDM` is built on the deprecated PyMC2 (***Patil et al., 2010***) backend, which notably is not yet built around an autodiff framework (***Paszke et al., 2019; Phan et al., 2019***), introduced with PyMC3 and beyond, rendering it increasingly difficult and worthwhile to install, maintain, and extend. The regression interface in `HDDM`, moreover, did not fully support proper hierarchical mixed-effect regression, rendering some natural model candidates untestable for experimentalists. While the LAN extension majorly expanded the reach of `HDDM` into broader model classes, the support is limited since the differentiability of the underlying neural networks could not really be exploited natively through `HDDM`.`HSSM` is designed as the direct successor, migrating the ecosystem to modern PyMC (***Abril-Pla et al., 2023***), adding JAX-based (***Phan et al., 2019***) hardware acceleration while preserving and extending the core capabilities that drove `HDDM`’s adoption.

### Tools based around Fokker Planck Equations

A few toolboxes have been developed with a focus on fast numerical solutions to solve Fokker Planck equations, which can in turn be leveraged to model a large collection of SSM variations, specifically enabling complex within trial dynamics to be incorporated. In particular the `PyDDM` (***Shinn et al., 2020***) and PyBEAM (***Murrow and Holmes, 2024***) libraries are based on this numerical foundation.

`PyDDM` is well-engineered, uses highly efficient numerical solvers (***Thomas, 2013***) and provides a useful GUI for model exploration. However, to the authors’ knowledge, `PyDDM` is restricted to two-choice diffusion models, does not currently support race models (***Heathcote and Matzke, 2022***), LBA variants (***Brown and Heathcote, 2008***), or n-choice architectures, and is moreover focused on maximum likelihood estimation rather than hierarchical Bayesian inference. It also does not natively support trial-by-trial neural regressors or hierarchical mixed-effect structures.

PyBEAM (***Murrow and Holmes, 2024***) extends the Fokker-Planck approach to a broader class of binary evidence accumulation models, supporting phenomena such as leaky integration, urgency signals, and time-varying evidence. It integrates with PyMC for Bayesian inference, however is also restricted to two-choice models, does not have an easy user-interface for hierarchical mixed-effects regressions, and does not provide built-in trial-by-trial neural covariates or domain-specific validation plots.

Its model space, while broader than the classical DDM, remains narrower than what simulation-based inference methods can accommodate and it wasn’t a priori designed for extension to those methods.

HSSM complements `PyDDM` by incorporating its numerical solvers as an alternative likelihood source within the `ssm-simulators` training data pipeline. Downstream the `HSSM` ecosystem then provides the hierarchical Bayesian inference layer that `PyDDM` lacks.

In other work, we explore some of the strength and weaknesses of Fokker Planck based approaches, in particular focusing on the implementation in PyDDM, to motivate fast analytical algorithms for a particular class of SSMs (***Liu et al., 2025, 2026***). In fact `PyDDM` (***Shinn et al., 2020***) specifically struggles with trial-wise definition of parameters, because this undercuts the free reuse of a once computed solution on a collection of trials, a driver of inference speed in the toolbox. The surrogate (***Fengler et al., 2021***), as well as fast analytical (***Navarro and Fuss, 2009***) likelihoods that we rely on in HSSM, benefit from batch computation and are inherently defined trial-wise. This enables fast likelihood computation in `HSSM`, regardless of the presence of trial-wise or solely global parameter specifications of the underlying cognitive process model.

### R Language

Beyond the Python ecosystem, several robust toolboxes have been developed in the R programming language. `EMC2` (***Stevenson et al., 2026***) is a particularly capable framework that supports hierarchical Bayesian estimation of multiple SSM families with a growing corpus of included models (as of this writing including the DDM, LBA, racing diffusion model (RDM), as well as lognormal race models (LNR)) `EMC2` uses Particle Metropolis within Gibbs (PMwG) sampling (***Kuhne et al., 2026***) and offers a linear modeling language for mapping model parameters to experimental designs. The framework supports custom user-defined likelihoods, flexible covariance structures (diagonal, blocked, or full multivariate normal) for modeling individual differences, and comprehensive model comparison via marginal likelihoods and Bayes factors. Its PMwG sampler is well-suited for posteriors with strong parameter correlations.

Notably, `EMC2` (***Stevenson et al., 2026***) includes a “trends” system that allows trial-by-trial covariates and enables the inclusion of built-in learning rules that can be used to combine with reinforcement learning paradigms.

For researchers working within the R ecosystem on analytically tractable SSMs, `EMC2` is arguably the most feature-complete option currently available.

However, while they share many similarities, the `EMC2` and`HSSM` packages differ quite fundamentally in their architectural philosophy, with consequences for extensibility and future-proofness. `EMC2` is designed as a self-contained package: sampler, likelihood functions, design language, and plotting infrastructure are all implemented internally. This benefits tight integration, but implies that improvements to `EMC2` must be driven by the `EMC2` community itself.

`HSSM`, by contrast, is designed as a node in a broader ecosystem of independently maintained packages. It delegates probabilistic programming to PyMC (***Abril-Pla et al., 2023***) and NumPyro (***Phan et al., 2019***), regression syntax to Bambi (***Capretto et al., 2022***), diagnostics to ArviZ (***Kumar et al., 2019***), auto-differentiation to JAX/PyTensor (***Phan et al., 2019; Team et al., 2016***), and likelihood storage to HuggingFace (***Jain, 2022***).`HSSM` then provides targeted functionality that points this integration of specialized packages in a direction that is particularly helpful for cognitive process modeling, but acts in large part as an orchestrator.`HSSM` therefore places itself to naturally inherit improvements from any of these upstream packages — for example, new samplers contributed to PyMC, NumPyro (***Phan et al., 2019***), or BlackJax (***Cabezas et al., 2024***) become immediately available — without requiring internal changes to`HSSM` itself. Another natural example are the continuous improvements to ArViZ, which benefit easy model comparison, exploratory analysis and Markov chain diagnostics (***Vehtari et al., 2021; Vats and Knudson, 2021; Sivula et al., 2025***).

This architectural difference has a direct bearing on the intractable-model problem that motivates the `HSSM` ecosystem. While `EMC2` could in principle incorporate surrogate likelihoods (e.g., via R interfaces to Keras or Torch), doing so would require substantial internal engineering because `EMC2`’s PMwG sampler is not built on a general-purpose probabilistic programming framework with native support for arbitrary differentiable log-probability functions.

We also mention hBayesDM (***Ahn et al., 2017***) which provides hierarchical Bayesian estimation across a curated library of reinforcement learning and decision-making tasks using Stan. It offers user-friendly interface and covers a broad range of task paradigms (e.g., Iowa Gambling Task, Go/NoGo, delay discounting). However, hBayesDM (***Ahn et al., 2017***) uses pre-compiled Stan models, making it difficult for users to add new models without Stan expertise. It is fundamentally focused on choice probabilities (e.g., via softmax) and provides only limited SSM support (standard DDM and LBA), without arbitrary mixed-effects regression on model parameters, trial-by-trial neural covariates, or simulation-based likelihood approximation.

### Joint RL-SSM Modeling

A major frontier in cognitive modeling is the simultaneous fitting of choices and continuous reaction time distributions while linking model parameters to trial-by-trial learning dynamics. The `RLSSM` package (***Fontanesi et al., 2019***) addresses this directly by combining reinforcement learning with sequential sampling models (DDM, RDM, LBA variants) using PyStan for hierarchical Bayesian estimation. However, rlssm relies on a fixed menu of pre-compiled Stan models with limited extensibility and depends on the now-deprecated PyStan 2 (***Van Hoey et al., 2013***). `HSSM`’s `RLSSM` class is designed to supersede this approach through a combinatorial framework: rather than pre-compiling each RL-SSM combination, `HSSM` allows users to dynamically pair arbitrary learning rules with any registered decision process model. The resulting `RLSSM` model then inherits the ability to specify trial-level regressions on any of the fundamental parameters of the combined model.

### Amortized Inference

`BayesFlow` (***Radev et al., 2020***) is a general-purpose library for amortized Bayesian inference using deep generative networks. `BayesFlow` (***Radev et al., 2020***) was originally designed to train networks that directly output posterior distributions, enabling near-instantaneous inference after an upfront training phase, but was recently extended to include some network architectures and learning algorithms which target likelihood surrogates. `BayesFlow` (***Radev et al., 2020***) is model-agnostic and has been applied across many scientific domains.

`HSSM` embraces integration with `BayesFlow` as a provider of high quality likelihood approximations. The package is focused more strongly on the learning algorithms than an exploratory scientific workflow via Bayesian modeling. We hence consider it upstream of packages like `HSSM` and, while important to mention as a powerful tool in the broader simulation based inference and scientific machine learning community (***Cranmer et al., 2020***), we consider it out of scope for a detailed direct comparison.

### Summary

The existing landscape is characterized by a tradeoff between flexibility and accessibility. Packages that support flexible model specification (`PyDDM`, `BayesFlow`) tend to lack hierarchical regression infrastructure, while those that provide hierarchical Bayesian estimation with formula syntax (`EMC2`, `hBayesDM`) rely on analytical likelihoods or fixed model libraries.`HSSM` is designed to resolve this tradeoff by combining an extensible, simulation-based approach to likelihood computation with a full-featured hierarchical regression interface, domain-specific validation tools, and a modular ecosystem architecture that encourages community contributions. To our knowledge, no existing package provides the combination of all features listed in Table 1

## Future Developments

The `HSSM` ecosystem is currently in beta status with well defined and tested core functionality. However, fundamentally, this establishes only a solid beginning with many directions for enhancements in vision.

First, we will incorporate more model classes. This includes models that intersect with our applied research initiatives, as well as models that have borne out general interest in the computational cognitive science community. Specifically, we are working on the inclusion of that embed within-trial dynamics which can vary according to other dynamic covariates such as moment-to-moment attention, a known special case of this framework is the attentional Drift Diffusion Model (***Krajbich et al., 2012***). We leverage recent developments concerning fast and accurate methods for estimating such models (***Liu et al., 2026, 2025***). Other candidates are circular diffusion models, as well as race models. Additionally, we are planning to cover a wider range of learning rules to be used through our `RLSSM` class, as well as more choice only models.

Second, we are planning to simplify our ability to draw approximate likelihoods from a variety of upstream sources. Our documentation includes a first example on how we can leverage the `BayesFlow` (***Radev et al., 2020***) as well as the `sbi` (***Tejero-Cantero et al., 2020***) packages to generate likelihood to be used downstream through HSSM. We are planning to deepen this integration and add other source packages for likelihood approximation such as `PyDDM` (***Shinn et al., 2020***), which is already incorporated as an alternative training data generator in the `ssm-simulators` package. A core piece to a smooth user-experience on this front will be the homogenization of our Hugging-Face repository across different likelihood sources.

Third, we are planning to leverage recent developments in the capabilities of large language models and corresponding agentic workflows. The modular design of the ecosystem — simple simulator and training data generation APIs, configurable likelihood constructors, and a high-level model-building syntax that maps cleanly to natural language descriptions of experimental hypotheses — makes its core components well-suited for emergent paradigms of workflow orchestration. Each step of a typical computational modeling pipeline (simulate from a candidate model, generate training data for a likelihood surrogate, construct a hierarchical regression, run inference, evaluate fit via posterior predictive checks, iterate) can be expressed as a self-contained function call with structured inputs and outputs. These design choices make programmatic orchestration feasible in a way that more monolithic or GUI-dependent toolboxes may struggle to support.

We are pursuing this vision along two complementary fronts. The first is didactic: improving the user experience of the ecosystem through LLM-assisted interaction. Our ecosystem already contains a self-improving question-answer bot, *HSSMeister*, which can answer user questions across all ecosystem packages. The second concerns the automation of the modeling workflow itself. We are developing *HSSMCortex*, a library that builds toward automated ecosystem extensions such as the integration of new cognitive models, and a collection of agent skills that help standardize the developer experience and harmonize contribution standards across the ecosystem. Beyond these internal initiatives, we see potential for more ambitious automation — an agent that, given a dataset and a set of candidate models, autonomously runs parameter recovery checks, fits competing models, compares them via information criteria, and surfaces the results as a structured report. Frameworks such as LangChain (***Chase, 2022***), are ideally suited for building super-structures for highly elaborate exploratory workflows which has the potential to severely reduce the tedium in iterating on small model adjustments, allowing researchers to focus on the bigger picture.

## Long-term maintenance of the ecosystem

The modular architecture of the ecosystem is deliberately built to absorb research progress on multiple fronts without requiring disruptive internal changes. At the simulation layer, the `ssm-simulators` package can be extended to incorporate alternative simulation backends: for example, using the Rust programming language as a backend (***Matsakis and Klock, 2014***) instead of our current Cython code (***Behnel et al., 2010***) for performance-critical models. Users will then be able to access the resulting simulators through the same high level interface that currently exists. At the training data layer, new generation mechanisms can be incorporated into the modular `TrainingData` class to serve the demands of alternative downstream libraries such as `BayesFlow` (***Radev et al., 2020***) and the sbi toolbox (***Tejero-Cantero et al., 2020***), broadening the set of surrogate likelihood sources available to the ecosystem. At the likelihood layer, `LANFactory` can be extended to accommodate novel neural network architectures; `HSSM`’s flexible likelihood constructors are designed to absorb these improvements automatically, both for inference and for fast prior and posterior predictive sampling via the linked simulators in `ssm-simulators`.

Our ecosystem interacts with a foundation of dependency packages that are continually enhanced. First, it depends on the PyMC (***Abril-Pla et al., 2023***) probabilistic programming library to construct computational graphs from model specifications. In the spirit of maintaining a high degree of flexibility, PyMC allows access to a host of different libraries which can be used for performing parameter inference on probabilistic models, including NumPyro (***Phan et al., 2019***) and BlackJax (***Cabezas et al., 2024***). Second, our toolkit depends on some auto-differentiation libraries, but again we maintain a high degree of flexibility, allowing any of Jax (***Frostig et al., 2019***), Pytorch (***Paszke et al., 2019***) as well as PyTensor (***Al-Rfou et al., 2016***) as backends. Together this set of dependencies allows us to benefit from advances in probabilistic programming, advances in deep learning as well as a solid foundation on the scientific python stack.

Building on PyMC (***Abril-Pla et al., 2023***) as the model construction foundation also opens the door to modular compositional classes that bring combinatorial modeling capabilities to end-users — an approach we are already exploring with our `RLSSM` class (combining decision processes with learning processes), and which can be extended in future work to e.g. GLM-HMM (Generalized Linear Model - Hidden Markov Model) frameworks or time-series approaches to lapse trial identification.

## Conclusions

A successor to the widely used but now outdated `HDDM` toolbox (***Wiecki et al., 2014***),`HSSM` distinguishes itself from existing alternatives (***Radev et al., 2020; Piray et al., 2019; Murrow and Holmes, 2024; Shinn et al., 2020; Stevenson et al., 2026***) through the combination of several properties that, to our knowledge, no single package currently provides: support for a large and growing bank of models with transparent likelihood management (analytic or surrogate), fully hierarchical between- and within-subject modeling with trial-wise neural regressors, gradient-based and GPU-accelerated inference, and a suite of domain-specific validation and visualization tools including posterior predictive checks, quantile probability plots, and model cartoon plots.

Critically, the ecosystem is designed not merely as a tool for end-users but as a platform for community contribution — the streamlined path from simulator to trained likelihood to registered `HSSM` model is intended to incentivize theoreticians to make their models available for empirical testing, creating a virtuous cycle between model development and experimental application. By providing a shared substrate with durable connective tissue, the ecosystem aims to reduce the reliance on project-specific, often unmaintained code-bases that currently still fragment our field and to broaden the range of theoretical models that experimental data can be tested against.

Built on the mature probabilistic programming foundation of PyMC (***Abril-Pla et al., 2023***) and Bambi (***Capretto et al., 2022***), the ecosystem is positioned to benefit from ongoing advances in computational statistics, deep learning, and simulation-based inference. Looking ahead, we plan to substantially expand the array of supported model classes, deepen integration with alternative likelihood sources, and explore the use of agentic workflows to automate and accelerate the iterative modeling process. We believe that the `HSSM` ecosystem can serve as the connective tissue between theoretical and experimental contributions to computational cognitive science — ultimately accelerating the research cycle from theory to empirical insight across both basic science and clinical applications.

## Acknowledgments

This work was supported by the National Institute of Mental Health (grants P50 MH119467-01 and P50 MH106435-06A1), the Office of Naval Research (MURI Award N00014-23-1-2792), and the Brain-storm Program at the Robert J. and Nancy D. Carney Institute for Brain Science. Moreover, this work was conducted using computational resources and services at the Center for Computation and Visualization, Brown University, which is supported by the National Institutes of Health (NIH) (Grant No. S10OD025181). We would moreover like to thank Jason Leng and Ivan Grahek for using the toolbox in earlier stages and providing invaluable user feedback as well as Isabella Aslarus, for her generous artistic contribution of our current set of package logos.

## Code and software availability

All packages described in this paper are released and are publicly available:

1. `HSSM`: https://github.com/lnccbrown/HSSM Documentation: https://lnccbrown.github.io/HSSM/ Installation: `pip install hssm`
2. `ssm-simulators`: https://github.com/lnccbrown/ssm-simulators Documentation: https://lnccbrown.github.io/ssm-simulators/ Installation: `pip install ssm-simulators`
3. `LANFactory`: https://github.com/lnccbrown/LANfactory Documentation: https://lnccbrown.github.io/LANfactory/ Installation: `pip install lanfactory`

The HuggingFace repository hosting trained likelihood networks is available at https://huggingface.co/franklab/HSSM.

## Use of AI tools in manuscript preparation

During preparation of this manuscript, the authors used Claude (Anthropic) as an editorial and writing assistant. This includes: feedback on draft structure and argumentation; refinement of prose for clarity and consistency; drafting initial versions of pedagogical content and figure captions, all subsequently reviewed, revised, and approved by the authors; assistance with L^A^TEX formatting and troubleshooting; and design assistance for schematic figures.

All scientific claims, model specifications, code, software implementations, citations, empirical examples, and analytical content presented in the paper were authored and verified by the human authors. No portion of the analysis, methodology, or findings was generated by AI. All AI-assisted draft material was reviewed and edited by the authors before inclusion.

AI tools were integrated in the later phases of the underlying software development, in line with emerging modern software engineering workflows.

## Supplementary figures

**Figure 3—figure supplement 1.**
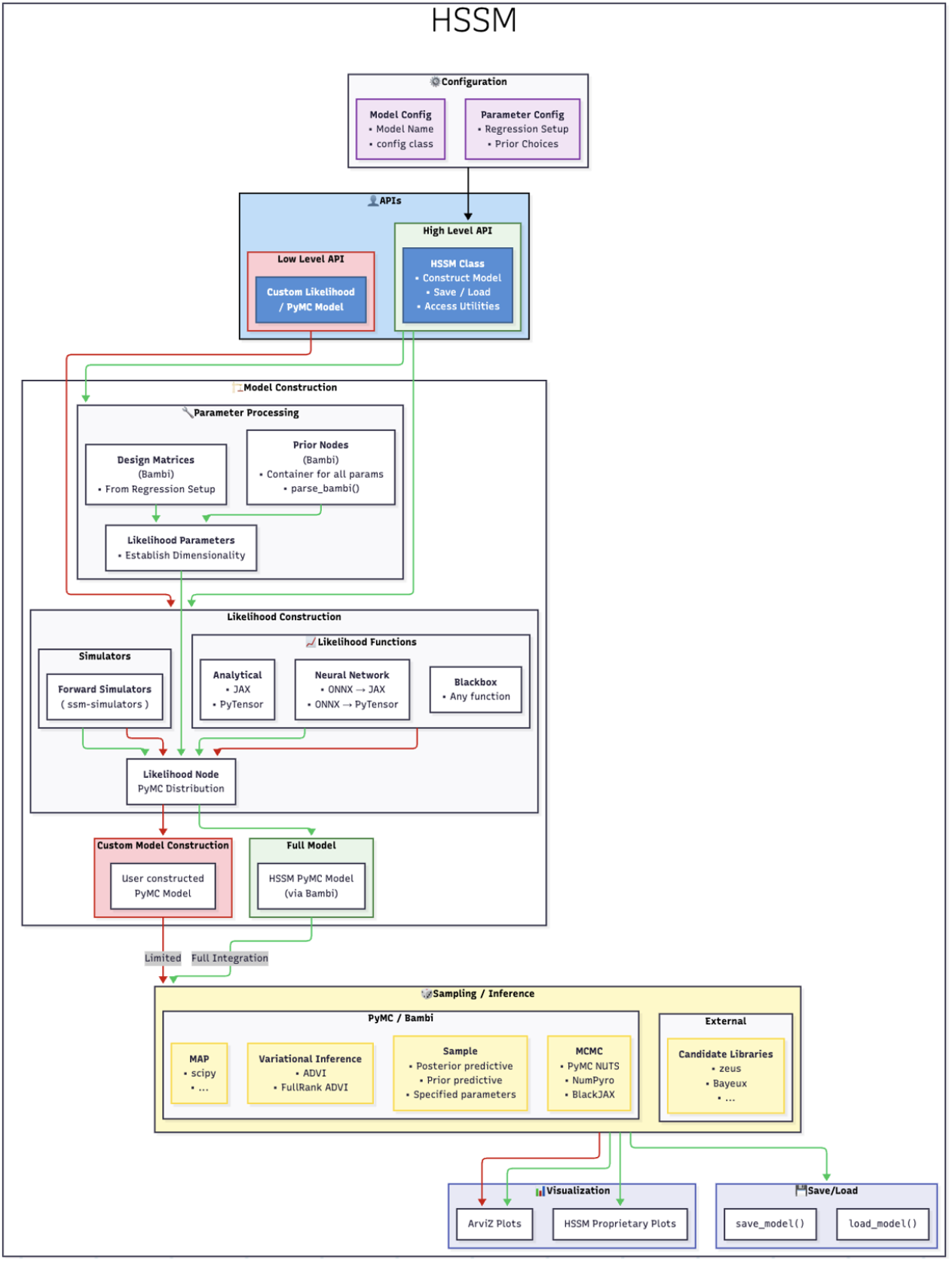
A more complete view on the architecture of the HSSM package. HSSM serves as the user-facing hub of the ecosystem and is designed around two complementary APIs (blue block). The high-level API centers on the `HSSM` class and consumes model and parameter configurations — including regression setup and optionally prior choices — to construct a full hierarchical Bayesian model automatically. The low-level API exposes the underlying pre-assembled PyMC distributions, allowing advanced users to build entirely custom PyMC models while retaining access to HSSM’s cognitive process model likelihoods. The likelihood layer accommodates multiple sources — analytical closed-form expressions, neural network surrogates loaded from ONNX (via either JAX or PyTensor), and blackbox user-provided functions — alongside forward simulators drawn from `ssm-simulators` for prior and posterior predictive sampling (users can provide their own if desired). The assembled likelihood node is a standard PyMC distribution and can be fit with a wide range of inference backends (yellow block): gradient-based MCMC via PyMC NUTS, NumPyro, or BlackJax; variational inference (ADVI, FullRank ADVI); MAP estimation; or external samplers. Outputs are compatible with ArviZ diagnostics and HSSM’s proprietary plotting utilities, and models can be persisted via native save/load methods.

**Figure 8—figure supplement 1.**
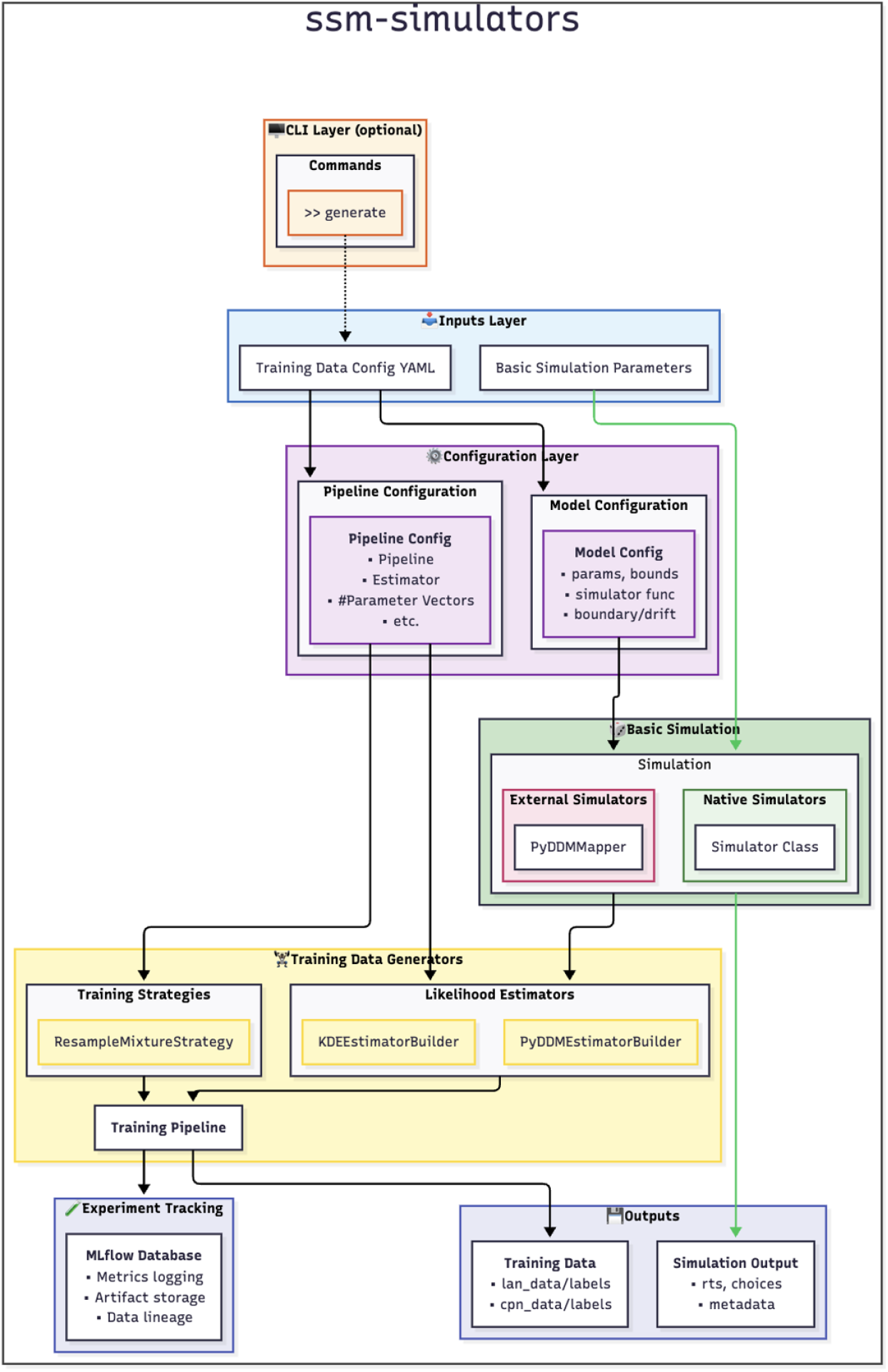
Architecture of the ssm-simulators package. The package provides two core functionalities: forward simulation of cognitive process models via the `Simulator` class, and a modular training data generation pipeline that combines configurable likelihood estimators and sampling strategies to produce structured datasets for downstream likelihood approximation. The package has a modular organization across level of functionality. Simulators can be constructed natively via `ssm-simulators` or e.g. via the `PyDDM` backend (***Shinn et al., 2020***) (with a vision toward extension). Training data generators are designed to combine different types of likelihood estimators as training signals for downstream networks (currently implemented are likelihoods based on `PyDDM` (***Shinn et al., 2020***) and our native approach following (***Fengler et al., 2021***).

**Figure 9—figure supplement 1.**
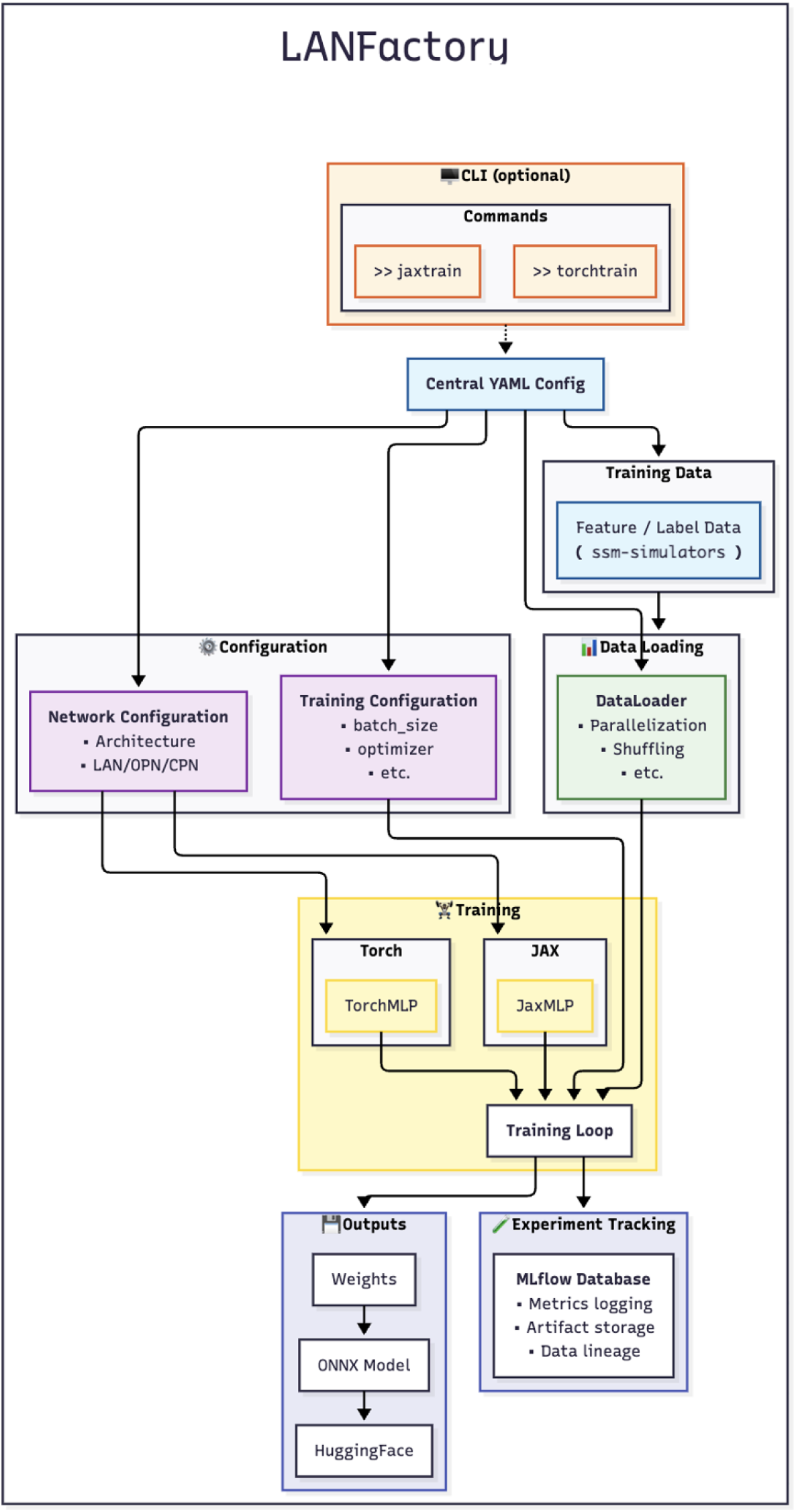
Detailed architecture of the LANFactory package. The package is configured through a central YAML file, optionally invoked via the `jaxtrain` and `torchtrain` CLI commands. The configuration drives three parallel substructures: network configuration (architecture choice and target network type — LAN, OPN, or CPN), training configuration (batch size, optimizer, and other hyperparameters), and data loading (parallelized, shuffled feature/label data, typically produced by `ssm-simulators`). The training subsystem provides parallel JAX and PyTorch implementations via the `JaxMLP` and `TorchMLP` class abstractions, both feeding into a unified training loop. Trained weights are exported to ONNX and uploaded to HuggingFace for downstream consumption by `HSSM` or other tools. Throughout training, an optional MLflow integration logs metrics, stores artifacts, and tracks data lineage to support reproducible network development.

## Notes

### Competing Interest Statement

The authors have declared no competing interest.

